# Microtubule attachment and centromeric tension shape the protein architecture of the human kinetochore

**DOI:** 10.1101/2020.02.11.944694

**Authors:** Alexander A. Kukreja, Sisira Kavuri, Ajit. P Joglekar

## Abstract

The nanoscale protein architecture of the kinetochore, a complex protein machine, plays an integral role in the molecular mechanisms underlying its functions in chromosome segregation. However, defining this architecture in human cells remains challenging because of the large size and compositional complexity of the kinetochore. Here, we use Förster Resonance Energy Transfer to reveal the architecture of individual kinetochore-microtubule attachments in human cells. We find that the microtubule-binding domains of the Ndc80 complex cluster at the microtubule plus-end. This clustering occurs only after microtubule attachment, and it increases proportionally with centromeric tension. Surprisingly, this clustering is independent of the organization and number of centromeric receptors for Ndc80. Moreover, Ndc80 clustering is similar in yeast and human kinetochores despite significant differences in their centromeric organizations. These and other data suggest that the microtubule-binding interface of the human kinetochore behaves like a flexible “lawn” despite being nucleated by repeating biochemical subunits.

## Introduction

To accurately segregate chromosomes, the kinetochore performs two different functions. When unattached, it acts as a signaling hub to delay the onset of anaphase, but when attached to the plus-ends of spindle microtubules, it acts as a force generating machine. The nanoscale organization of kinetochore proteins relative to one another and relative to the microtubule plus-end, referred to here as the “architecture” of the kinetochore, plays key roles in the molecular mechanisms underlying both of these functions (Aravamudhan et al., 2014; Aravamudhan et al., 2015; McIntosh, 1991; Wan et al., 2009). However, the architecture of the human kinetochore has not yet been defined because its features make it intractable to many forms of microscopy. The human kinetochore is compositionally complex and large, built from hundreds of protein molecules distributed upon a 200 nm diameter disk-like chromatin foundation known as the centromere. Furthermore, it changes in response to microtubule attachment and force (Magidson et al., 2016; Magidson et al., 2015; Suzuki et al., 2014).

Because no one method can currently define kinetochore architecture, it must be synthesized from data defining four fundamental aspects: (1) the structures of kinetochore proteins (see (Hinshaw and Harrison, 2018; Musacchio and Desai, 2017)), (2) their copy numbers, (3) their average localizations along the kinetochore-microtubule axis, and (4) their nanoscale distribution around and along the plus-end (Joglekar and Kukreja, 2017). For the human kinetochore, data regarding the first three aspects are available (Magidson et al., 2016; Smith et al., 2016; Suzuki et al., 2015; Suzuki et al., 2014; Suzuki et al., 2018; Wan et al., 2009). However, the nanoscale distributions of kinetochore proteins relative to microtubule plus-ends remains unknown. Here we apply Förster Resonance Energy Transfer (FRET) between fluorescently labeled kinetochore subunits to elucidate this aspect of the human kinetochore.

We designed our FRET quantification experiments to elucidate specific aspects of the architecture of the human kinetochore. Our primary goal was to determine the organization of the microtubule-binding Ndc80 complex (Ndc80C) relative to the microtubule plus-end, because Ndc80C is necessary for forming end-on microtubule attachment, force generation, and it can regulate plus-end polymerization dynamics (Ciferri et al., 2008; DeLuca et al., 2006). The human kinetochore contains ∼ 250 molecules of Ndc80C and binds ∼ 20 microtubule plus-ends, suggesting that ∼ 12 Ndc80C molecules engage one microtubule plus-end (Suzuki et al., 2015; Wendell et al., 1993). The nanoscale distribution of these molecules around the 25 nm diameter and along the longitudinal axis of the microtubule will influence the persistence of the kinetochore-microtubule attachment (Hill, 1985). However, the distribution of Ndc80C molecules is dictated by the long and flexible centromere-bound protein linkages. Therefore, we extended our FRET analysis to members of the <u>C</u>onstitutive <u>C</u>entromere-<u>A</u>ssociated <u>N</u>etwork (CCAN) of proteins involved in Ndc80C recruitment. Microtubule attachment- and tension-dependent changes in kinetochore architecture are at the heart of the its ability to implement emergent functions. Therefore, we also studied how the nanoscale distributions of kinetochore components change in response to attachment, tension, and perturbation of proteins that link it to the centromere. Finally, we use our FRET data to formulate a model of a human kinetochore-microtubule attachment and contrast it with the yeast kinetochore.

## Results

### Design and implementation of a FRET imaging strategy to study kinetochore architecture

To determine protein proximities in HeLa kinetochores using FRET, we co-expressed EGFP (donor fluorophore, referred to as GFP) and mCherry (acceptor fluorophore) labeled kinetochore proteins (see Methods; (Khandelia et al., 2011)). To maximize the recruitment of the labeled proteins to the kinetochore, we also knocked down their endogenous, unlabeled counterparts using RNAi (Fig. S1). Depending on the position of the donor and acceptor fluorophores (fused to either the C- or N-terminus of the selected proteins), we expected FRET to occur within a single protein complex (intra-complex), between neighboring complexes (inter-complex, left and middle panel in Fig. 1A), or both (right panel in Fig. 1A).

**Figure 1.**
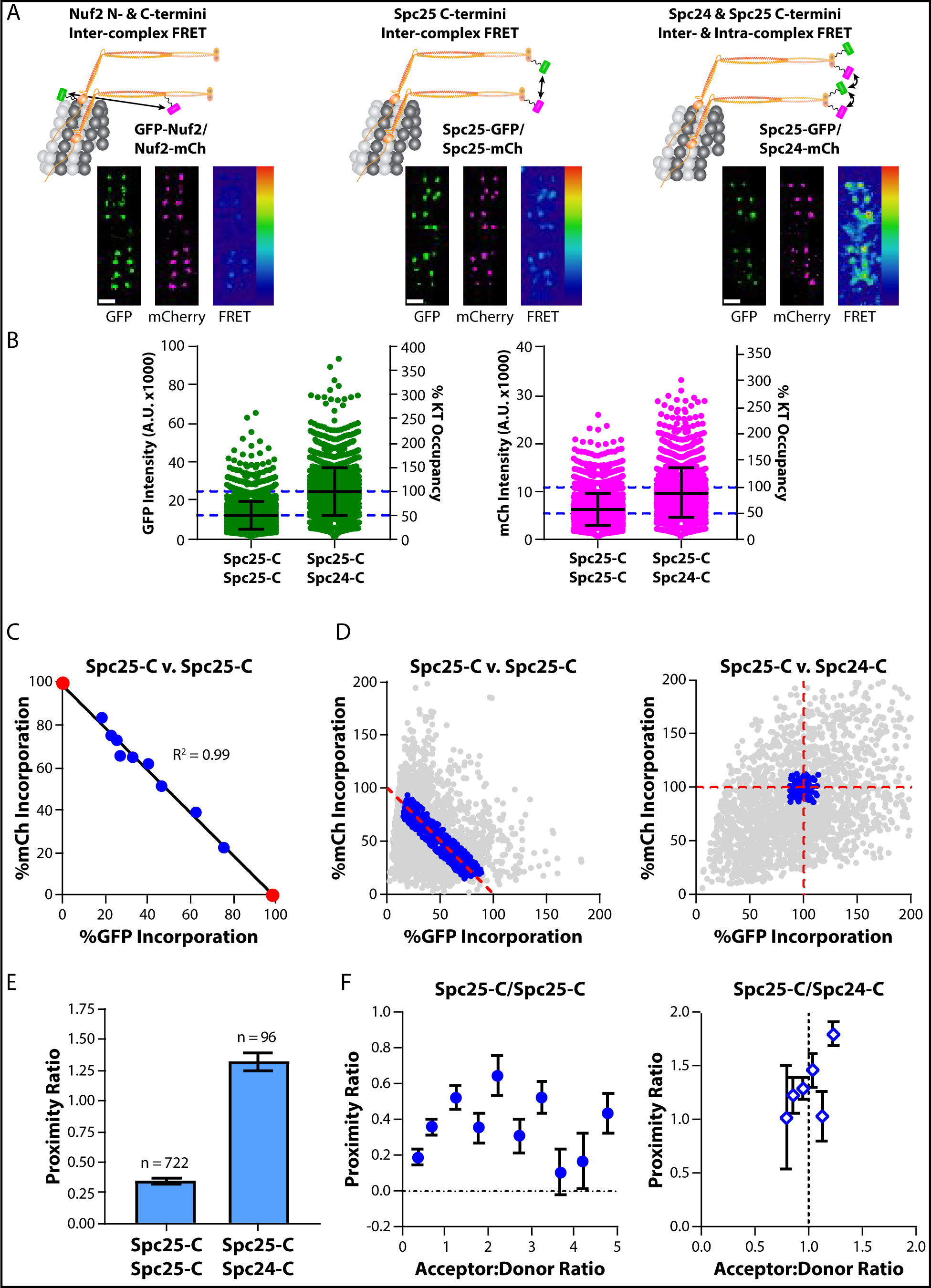
Construction of a FRET quantification methodology to determine protein proximities at human kinetochores. A) Top: Co-expression of GFP (FRET donor, green) and mCherry (mCh, FRET acceptor, magenta) fusions of the Ndc80 complex (Ndc80C, orange) reveal the proximity between adjacent Ndc80C molecules along the longitudinal axis (left), around the circumference of the microtubule (middle, right). For simplicity, the cartoons depict only two Ndc80C molecules per microtubule. Micrographs show representative metaphase plates from each cell line. FRET micrographs are scaled equivalently; GFP and mCherry micrographs are scaled for ease of viewing. Scale bar, 1 µm B) GFP and mCherry fluorescence intensity of individual kinetochores in cells expressing Spc25-GFP/Spc25-mCherry and Spc25-GFP/Spc24-mCherry after siRNA-mediated knockdown of endogenous, unlabeled Spc25 or both Spc25 and Spc24, respectively. The Y axis on the right shows the saturation level of the kinetochore by the GFP- and mCherry labeled subunit, estimated from (Suzuki et al., 2015) C) Correlation between Spc25-GFP and Spc25-mCherry signals measured from kinetochores from the data set in B. Data were binned by the ratio of mCherry to GFP fluorescence, and further normalized by the X- and Y-intercepts of their linear regression (black line; see main text). Measurements of cells expressing Spc25-GFP or Spc25-mCherry in isolation are marked by red circles. Bin sizes, from left to right: n = 398, 379, 109, 108, 131, 145, 170, 212, 491, 320, 499. D) Normalized fluorescence intensity for all kinetochores measured in Spc25-C/Spc25-C (left) and Spc25-C/Spc24-C (right) expressing cells. Only the data indicating complete saturation of the kinetochore by fluorophore-labeled proteins (blue circles) were used to measure FRET. All other values are excluded (gray circles) E) The proximity ratio for fully-occupied, metaphase kinetochores in Spc25-C/Spc25-C and in Spc25-C/Spc24-C expressing cells. The number of kinetochores measured for each cell line is indicated above the bars. F) Dependence of the proximity ratio on the ratio of the number of acceptor-to donor-labeled molecules (A:D). In the absence of competition, the proximity ratio clusters around an A:D that reflects the inherent stoichiometry of the two fluorophore-labeled subunits involved (Spc25-C/Spc24-C cells, right). Data are binned by A:D (mean ± SEM; for Spc25-C/Spc25-C, n = 150, 192, 93, 62, 37, 53, 50, 36, 24, 25; for Spc25-C/Sp24-C, n = 2, 13, 32, 30, 14, 5). In B), error bars are ± S.D. In C), E), and F) data represent the mean ± SEM In C), SEM error bars are too small to be seen. Data collected in B) – F) are from N ≥ 3 experiments.

For FRET to accurately reveal protein proximities, kinetochores must be saturated by the donor- and acceptor-labeled proteins. However, in cells co-expressing GFP- and mCherry-labeled versions of Spc25, a Ndc80C subunit, we observed significant variability in kinetochore signals, indicating that kinetochores in many cells were not fully saturated with labeled proteins (Fig. S2). We expected the variability to mainly arise from changes in fluorescence intensity that occur with depth from the coverslip, cell-to-cell variation in siRNA efficiency, and to a smaller extent, overlapping signals from neighboring kinetochores. To minimize the effects of this variability and measure FRET only from kinetochores fully saturated with labeled proteins, we established an objective filtering scheme as follows.

We quantified the GFP and mCherry fluorescence signals per kinetochore in cells co-expressing Spc25-GFP and Spc25-mCherry (Spc25 is a subunit of Ndc80C, Fig. 1B). Because the kinetochore has a limited number of protein binding sites, we expected the donor- and acceptor-labeled versions of Spc25 to compete for kinetochore binding. To reveal this competition, we minimized the noise in the fluorescence data by binning the signals from each kinetochore by the ratio of the mCherry to GFP fluorescence intensity. The mean values of binned data showed a clear anti-correlation between the Spc25-GFP and Spc25-mCherry signal per kinetochore (Fig. 1C, blue circles). We performed linear regression of the data to determine the X- and Y-intercepts (Fig. 1C and Fig. S3A; note that the data in figure 1C and 1D have been normalized by the X- and Y-intercepts of the linear regression). These intercepts should correspond to intensities of kinetochores fully saturated with GFP and mCherry respectively. Accordingly, the average fluorescence intensity per kinetochore in cells that expressed either Spc25-GFP or Spc25-mCherry alone were in agreement with the respective intercept values (Fig. 1C, red circles). Therefore, we used the X- and Y-intercept values with the known copy number for Ndc80C molecules per kinetochore to define the single molecule brightness of GFP and mCherry (see Methods; (Suzuki et al., 2015)). Using these single molecule brightness values, we converted the GFP and mCherry fluorescence intensities from each kinetochore into protein counts and retained only the measurements reflecting full kinetochore occupancy (Fig. 1D, blue circles).

To quantify FRET, we determined the acceptor fluorescence due to FRET, which is known as ‘sensitized emission’. The sensitized emission value for each kinetochore was calculated by subtracting the contributions of GFP bleed-through and mCherry cross-excitation from the signal measured in the FRET channel (Fig. S3, Methods). Because the sensitized emission is directly proportional to the average FRET efficiency and the total number of FRET pairs, we normalized it with respect to the number of donor and acceptor molecules per kinetochore. This normalization renders a FRET metric referred to as the ‘proximity ratio’. The proximity ratio is expected to be proportional to the average FRET efficiency (Joglekar et al., 2013; Muller et al., 2005).

Using this methodology, we measured a moderate proximity ratio between the C-termini of Spc25 subunits and a high proximity ratio between the C-termini of Spc25 and Spc24 (Fig. 1E; we will refer to FRET between fluorophores as FRET between the protein termini to which the fluorophores are fused; in this case, FRET between Spc25-C/Spc25-C and Spc25-C/Spc24-C, respectively). Thus, the average inter-complex distance between neighboring Ndc80C molecules at their centromeric ends (as measured by Spc25-C/Spc25-C FRET) is < 10 nm. The much higher FRET measured between neighboring Spc25 and Spc24 molecules indicates that these C-termini are more densely organized than Spc25-C/Spc25-C. This is consistent with the intra-complex separation of ∼ 2 nm between Spc25-C and Spc24-C (Ciferri et al., 2008). It is worth noting that the competition between donor- and acceptor-labeled Spc25 molecules in Spc25-C/Spc25-C expressing cells will yield kinetochores with varying acceptor to donor ratios (A:D). This effect introduces variation in the measured proximity ratio at a given kinetochore. This effect is absent from the Spc25-C/Spc24-C FRET pair because these proteins do not compete with each other for kinetochore binding (Fig. 1F). Accounting for A:D, however, does not significantly change the trends and interpretation of our FRET data (see Supplementary Table 1). Therefore, all figures presented in this study display the average proximity ratio irrespective of A:D.

### Ndc80C molecules are clustered along their entire length and staggered along the longitudinal axis of the kinetochore-microtubule attachment

The nanoscale distribution of Ndc80C molecules relative to the microtubule plus-end governs the ability of the kinetochore to persistently attach to the plus-end and influence the polymerization dynamics of the microtubule (DeLuca et al., 2006; Hill, 1985). Therefore, to understand how adjacent Ndc80C molecules are aligned relative to one another, we measured inter-complex FRET by positioning fluorophores along the entire length of the Ndc80C. We chose three locations along the length: proximal to its microtubule-binding end (fused to N-Nuf2, wherein N-denotes the N-terminus of the protein), near the middle of its ∼ 57 nm span (fused to Nuf2-C which is proximal to its tetramerization domain), or near its centromeric end (fused to Spc25-C). We detected FRET at all three positions, indicating that the average distance between adjacent Ndc80C molecules is < 10 nm along its entire length. Furthermore, the proximity ratio was higher at the microtubule-binding end and in the middle of Ndc80C than at its centromeric end (Fig. 2A). Therefore, Ndc80C molecules are more tightly clustered on the microtubule lattice and at their tetramerization domains than at the ends which anchor them to the centromere.

**Figure 2.**
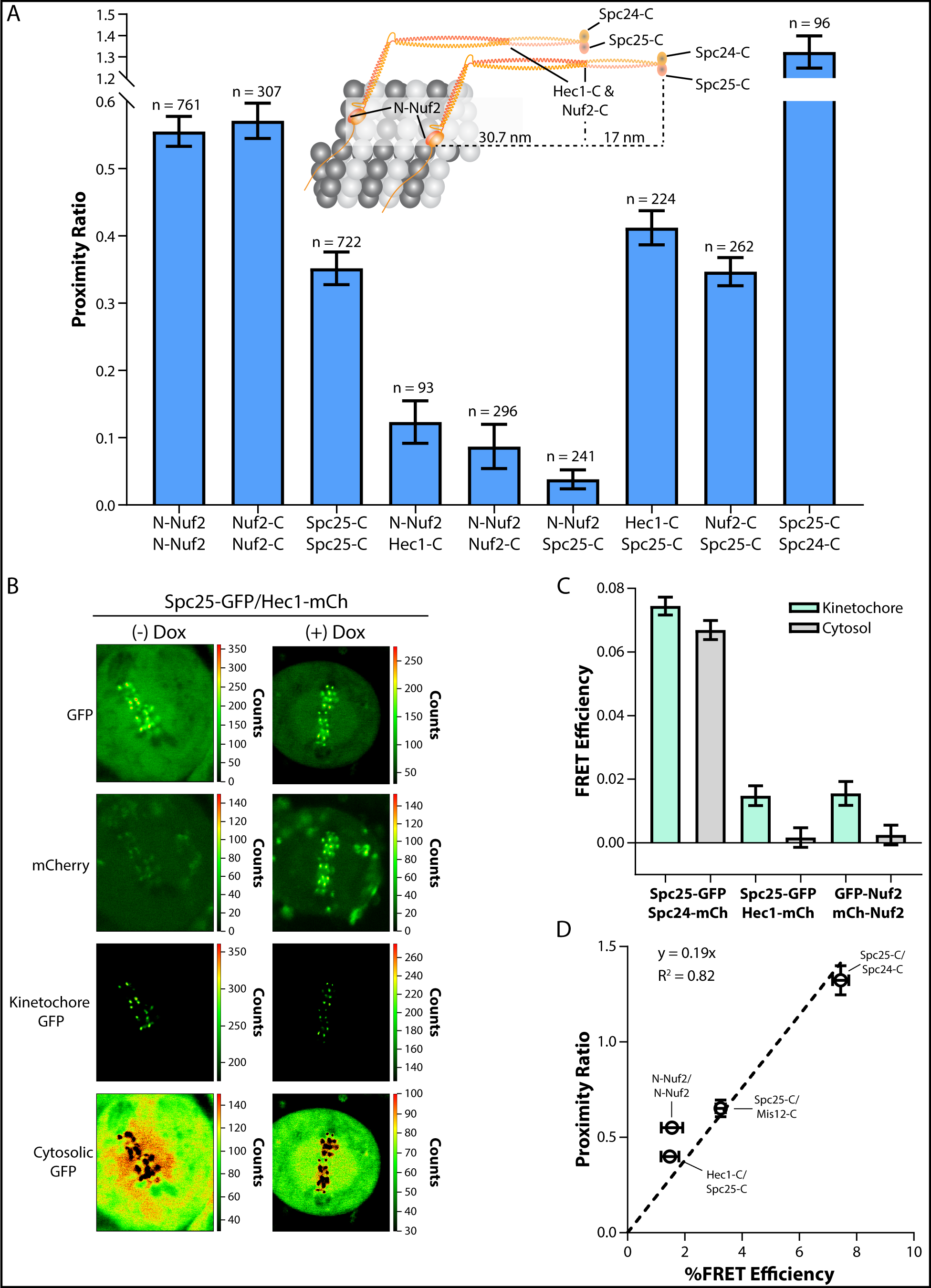
Ndc80C molecules are clustered along their entire length and staggered along the microtubule lattice in metaphase kinetochores. A) FRET measurements for subunits of Ndc80C in microtubule-attached metaphase kinetochores. Cartoon depicts the position and approximate distance between different anchoring points for donor and acceptor fluorophores. For simplicity, only two Ndc80C molecules are shown to illustrate how Ndc80C molecules may cluster and stagger when bound to the microtubule lattice. Bar graph displays the average proximity ratio ± SEM. The number of kinetochores measured for each cell line is indicated above the bars. B) Example confocal micrographs of Spc25-GFP/Hec1-mCherry HeLa cells for FLIM. Doxycycline (Dox) induces the expression of Hec1-mCherry. All images are scaled by the number of photons/pixel (scale to the right of images). Intensity thresholding was used to separate kinetochore-localized from cytosolic GFP pixels (bottom two rows). Note that in the absence of doxycycline, the GFP channel shows minor bleed-through into the mCherry channel. C) FRET efficiency of kinetochore-localized (light green bars) and cytosolic (gray bars) FRET pairs in the indicated cell lines. Bars represent the average FRET efficiency ± SEM. The number of measurements from each cell line (left to right) = 15, 14, and 13. D) Plot of the average proximity ratio versus the average FRET efficiency for each of the indicated FRET pairs fit with a linear regression. Error bars are ± SEM. Note that data from the N-Nuf2/N-Nuf2 expressing cell line deviates from the trend, potentially due to complications with controlling for the acceptor-to-donor ratio with the FLIM setup. All data are from N ≥ 3 experiments.

Because Ndc80C and the long, multivalent protein linkages that recruit it to the centromere are both flexible, it is possible that Ndc80C molecules can cluster without aligning with neighboring molecules along the long axis of the kinetochore-microtubule attachment (Ciferri et al., 2008; Wang et al., 2008). To reveal the extent of alignment between adjacent Ndc80C molecules, we next positioned fluorophores on two different subunits of Ndc80C (Aravamudhan et al., 2014; Joglekar et al., 2009; Wang et al., 2008). We chose fluorophore positions such that intra-complex should not occur; it can only arise between neighboring Ndc80C molecules that are staggered relative to each other along the longitudinal axis of the kinetochore-microtubule attachment. We did not detect FRET between fluorophores placed at the two extremes of Ndc80C, located ∼ 47 nm apart within a single complex (N-Nuf2/Spc25-C, Fig. 2A). We detected weak FRET, however, between fluorophores placed at N-Nuf2/Hec1-C (∼ 30 nm intra-molecular separation) and moderate FRET between Hec1-C/Spc25-C (∼ 17 nm intra-molecular separation). These measurements were further confirmed by similar proximity ratio values measured for FRET between N-Nuf2/Nuf2-C and Nuf2-C/Spc25-C (Fig. 2A). Thus, a measurable fraction of neighboring Ndc80C molecules are staggered relative to one another along the longitudinal axis of the microtubule. The average offset between adjacent molecules is < 30 nm, because minimal inter-complex FRET was detected between N-Nuf2/Hec1-C. This organization is significantly different from the organization of the budding yeast kinetochore, wherein adjacent Ndc80C molecules are aligned with each other (Aravamudhan et al., 2014).

### Fluorescence Lifetime Imaging confirms staggering of adjacent Ndc80C molecules

The conclusion that Ndc80C molecules are staggered along the microtubule lattice assumes that the detected FRET takes place between adjacent complexes. To independently confirm this, we measured FRET between Ndc80C molecules localized to the kinetochore and between molecules freely diffusing in the cytosol. If the observed FRET arises due to the arrangement of adjacent Ndc80C molecules in the kinetochore, then it should be detected only within kinetochores, but not in the cytosol. Conversely, if FRET occurs intra-complex, then it should be detectable both at the kinetochore and within the cytosol. We used Fluorescence Lifetime IMaging (FLIM) to simultaneously measure FRET in both populations of kinetochore proteins. FLIM directly measures FRET efficiency from the decrease in the excited-state lifetime of the donor fluorophore due to the presence of an acceptor within 10 nm (Becker, 2012). Since the fluorescence lifetime of the donor can be determined accurately even at low fluorophore concentration, we could separately quantify FRET between kinetochore-localized and cytosolic Ndc80C molecules.

We first tested the validity of this approach by measuring the efficiency of FRET between N-Nuf2/N-Nuf2 and between Spc25-C/Spc24-C. In the former case, only inter-complex FRET is possible, and hence it should be detected only within kinetochores. In the latter case, FRET is predominantly intra-complex, and it should occur within kinetochores and in the cytosol (Ciferri et al., 2008). We used an empirical intensity threshold to separate pixels corresponding to the kinetochores from the cytosol (Fig. 2B, S4, and Methods). Fluorescence lifetime measurements on these two classes of pixels confirmed our expectations. In cells expressing GFP-Nuf2/mCherry-Nuf2, the GFP fluorescence lifetime was lower in the kinetochore, but unchanged in the cytosol as compared to its lifetime in cells expressing GFP-Nuf2 alone (Fig. 2C). In cells expressing Spc25-GFP/Spc24-mCherry, the GFP lifetime was significantly lower in both kinetochores and the cytosol when compared to cells expressing Spc25-GFP alone. We next measured FRET efficiency between Hec1-mCherry and Spc25-GFP using FLIM. For the Hec1-C/Spc25-C pair, FRET was detectable only within kinetochores and not within the cytosol (Fig. 2C). Thus, intra-complex FRET does not occur with the Hec1-C/Spc25-C pair. These data further support the conclusion that adjacent Ndc80C molecules are staggered along the longitudinal axis of the kinetochore-microtubule attachment in human kinetochores.

It is worth noting that the FRET efficiencies measured from the FLIM experiments were directly proportional to the independently determined proximity ratio values (Fig. 2D). Thus, normalizing sensitized emission by the number of donor and acceptor molecules (i.e., the proximity ratio) reflects the average proximity between adjacent kinetochore subunits.

### The protein linkages that tether Ndc80C to the centromere are sparsely distributed

The clustered and staggered organization of Ndc80C molecules in attached kinetochores can result from the spatial organization of the protein linkages that tether Ndc80C to the centromere. In the human kinetochore, Ndc80C is recruited to the centromere via two different centromeric proteins: CenpC and CenpT (Fig. 3A, (Gascoigne et al., 2011; Huis In ’t Veld et al., 2016; Klare et al., 2015; Nishino et al., 2013; Rago et al., 2015; Screpanti et al., 2011; Weir et al., 2016)). CenpC and CenpT bind to the centromere using their C-terminal domains and extend their N-terminal domains outward to bind one Mis12 complex each (Mis12C). Mis12C is a ∼20 nm long linker/adaptor that in turn binds one Ndc80C. Additionally, the N-terminal domain of CenpT directly recruits two Ndc80C molecules (Huis In ’t Veld et al., 2016). Therefore, we next used FRET to elucidate the nanoscale organization of these linkages.

**Figure 3.**
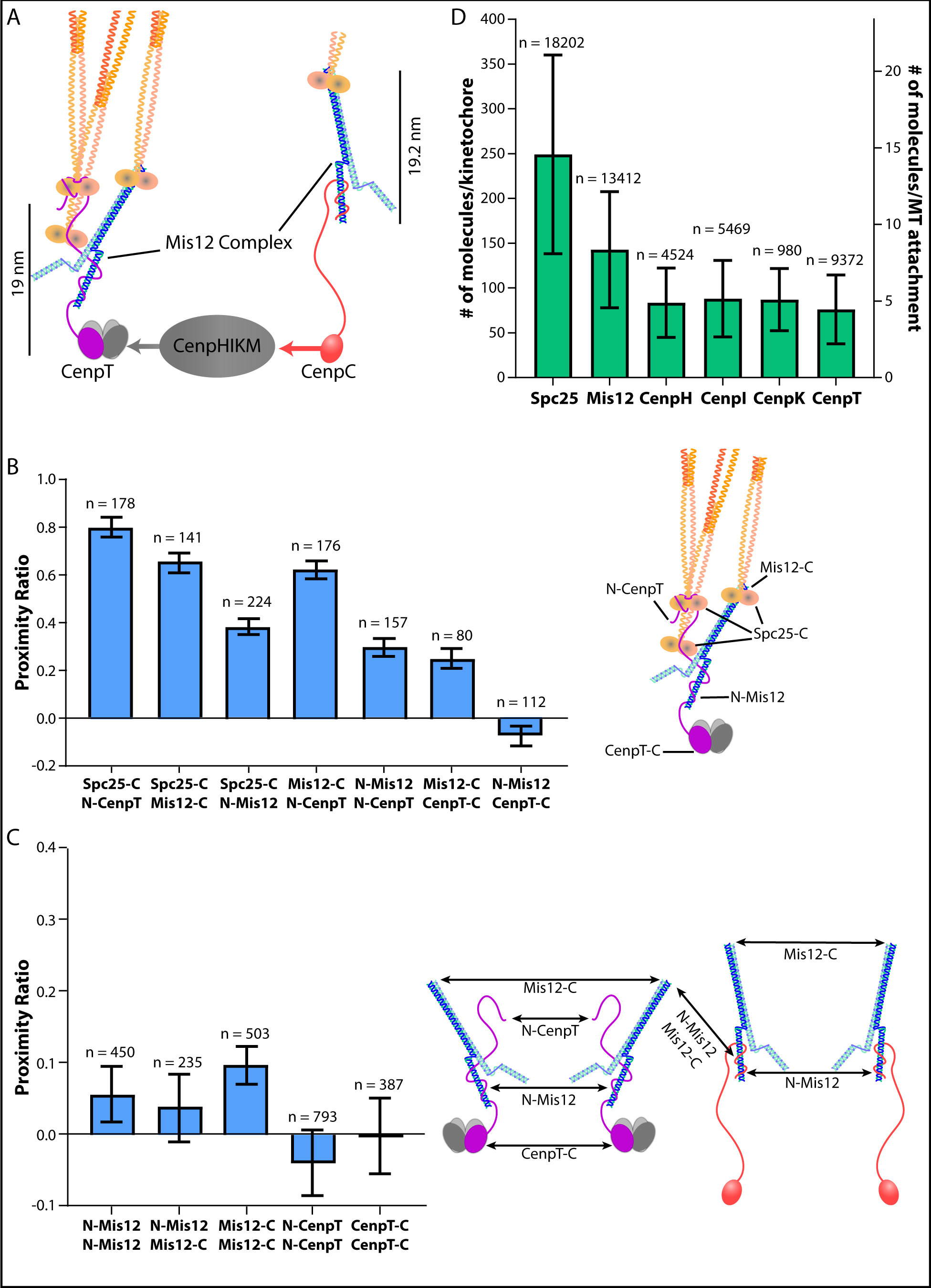
The CenpC and CenpT recruitment domains are ≥ 10 nm apart. A) Diagram of the biochemical recruitment pathway for Ndc80C. Only the half of Ndc80C which binds to the centromere is shown for simplicity. B) FRET between proteins involved in Ndc80C recruitment. Diagram to the right shows the location of different fluorophore tags within the CenpT linkage. Note that FRET between Ndc80C and Mis12C is also possible in the CenpC linkage C) The lack of FRET between neighboring Mis12C molecules and between neighboring CenpT molecules suggests that Ndc80C linkages are ≥ 10 nm apart. Diagram shows potential sources of FRET between individual CenpC and CenpT linkages. D) Protein copy numbers for metaphase kinetochores, evaluated from unfiltered fluorescence intensities of kinetochores in cells expressing GFP and/or mCherry labeled versions of the indicated subunits. Bar graphs in B) and C) display the average proximity ratio ± SEM. of fully occupied, metaphase kinetochores. Bar graph in D) displays the average number of molecules per kinetochore (left axis) or per microtubule (right axis, assuming 17.1 microtubules per human kinetochore) ± SD. The number of kinetochores measured for each cell line is indicated above the bars in B), C), and D). All data are from N ≥ 3 experiments

We first characterized the organization of proteins within the CenpT-Mis12C-Ndc80C linkage using FRET pairs positioned near sites of protein-protein interactions (Fig. 3B). The protein proximities revealed by these FRET measurements are consistent with the known organization of this linkage (Fig. S6B, (Huis In ’t Veld et al., 2016; Nishino et al., 2013; Petrovic et al., 2016). Next, we determined the proximity between neighboring Mis12C molecules. Mis12C serves as a convenient proxy for CenpC and CenpT distribution at the centromere, because both CenpC and CenpT bind only one Mis12C. We detected only weak inter-complex FRET between adjacent Mis12C molecules, irrespective of whether fluorophores were placed at N-Mis12, Mis12-C (Mis12 is a subunit of the Mis12 complex), and even when N- and C-terminal tagged Mis12 molecules were co-expressed (Fig. 3C). Thus, adjacent Mis12C molecules are, on average, ≥ 10 nm apart. We next measured inter-complex FRET between adjacent CenpT molecules. We did not detect FRET between the N- or the C-termini of CenpT molecules (Fig. 3C). Thus, neighboring CenpT molecules are also spaced ≥ 10 nm apart. We did not include CenpC in these analyses due to technical difficulties. However, the absence of inter-complex FRET between Mis12C molecules indicates that the domains of CenpC that recruit Ndc80C are also ≥ 10 nm apart.

An additional component that may influence the organization of Ndc80C is the CenpH, CenpI, CenpK, and CenpM (CenpHIKM) complex, a CCAN component (Basilico et al., 2014; Klare et al., 2015; McKinley et al., 2015). CenpHIKM organizes centromeric chromatin and bridges CenpT with CenpC (Fig. 3A). Consistent with this role, we found a ∼ 1:1 stoichiometry between three of the four CenpHIKM subunits with CenpT (average for CenpH, CenpI, and CenpK = 86.3 ± 0.8 (S.E.M.) molecules) from quantification of fluorescent protein intensities at kinetochores (Fig. 3D, (Suzuki et al., 2015)). FRET measurements between neighboring CenpI subunits revealed that, like CenpT and Mis12C, these subunits are also ≥ 10 nm apart (Fig. S5B). Finally, the protein termini of most of the CenpHIKM subunits were found to be proximal to the centromere and not within proximity of Ndc80C (Fig. S5).

In sum, these data reveal that adjacent protein linkages recruiting Ndc80C are ≥ 10 nm apart from one another. Nevertheless, Ndc80C molecules are clustered and staggered along the microtubule lattice. Two factors may contribute to this Ndc80C organization. First, the multivalent recruitment of Ndc80C molecules by CenpT could place multiple Ndc80C molecules within 10 nm, allowing for both clustering and staggering. Second, the span between the microtubule-binding domains of Ndc80C and the N-terminal domains of CenpT and CenpC is ∼ 50 – 70 nm. Thus, even though the CenpT and CenpC recruitment domains are not within FRET proximity, Ndc80C molecules recruited by distantly spaced molecules may still bind near each other to the same plus-end.

### Microtubule attachment clusters Ndc80C in both human and budding yeast kinetochores

We examined the role of microtubule-binding in Ndc80C organization by quantifying FRET between adjacent Ndc80C molecules in unattached kinetochores. For this, we destroyed the mitotic spindle in HeLa cells by treating them with nocodazole, a microtubule depolymerizing drug (Fig. 4A & S6). In unattached kinetochores, inter-complex FRET between adjacent Ndc80C molecules reduced significantly. The strongest decrease occurred at the microtubule-binding end (N-Nuf2), with a smaller decrease near the tetramerization domain (Nuf2-C), and only a modest reduction at its centromeric end (Spc25-C, Fig. 4B). The reduced FRET is unlikely to arise from structural rearrangement within Ndc80C, because FRET between Spc25-C and Spc24-C showed only a modest decrease (∼ 10%, likely reflecting reduced inter-complex FRET, Fig. 4B). Thus, binding to the microtubule plus-end is the main reason for the clustering of the microtubule-binding domains of Ndc80C in metaphase kinetochores.

**Figure 4.**
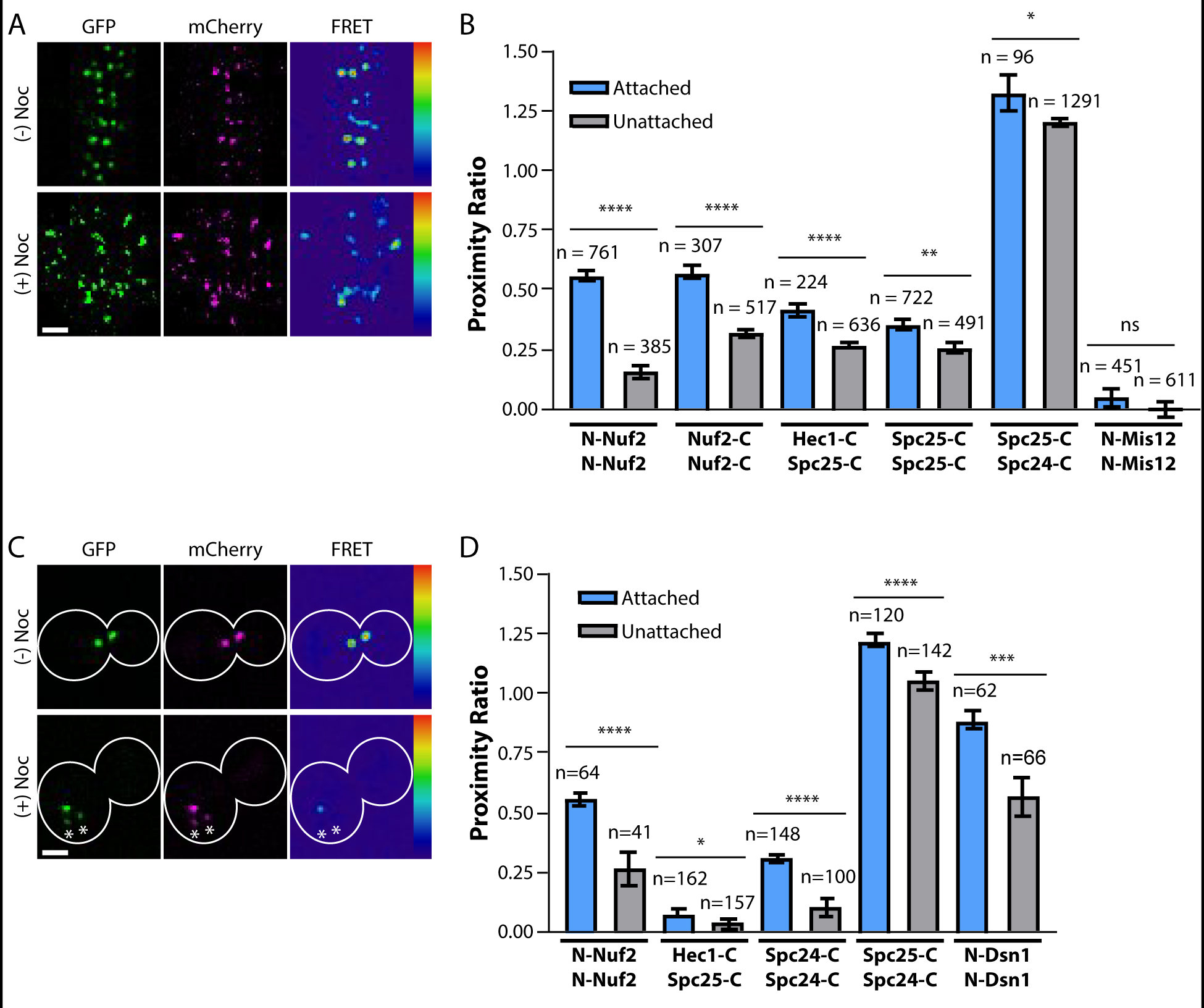
The spatial organization of Ndc80C is responsive to microtubule binding in both human and budding yeast kinetochores. A) Example micrographs of metaphase HeLa cells expressing N-Nuf2/N-Nuf2, with or without the addition of nocodazole (Noc). B) The lack of microtubule attachments reduces FRET between subunits of Ndc80C. Light blue bars are measurements from metaphase aligned kinetochores and gray bars are from nocodazole-treated, unaligned kinetochores. C) Example micrographs of budding yeast metaphase cells expressing N-Nuf2/N-Nuf2, with or without the addition of nocodazole. White outlines indicate the cell membrane. Asterisks highlight the position of small, dim clusters of unattached kinetochores (see Fig. S7). D) Same as in B) but as measured for budding yeast kinetochores. For A) and C), FRET micrographs are scaled equivalently; GFP and mCherry micrographs are scaled for ease of viewing. Scale bar, 2 µm. For D) and B), bar graphs display the average proximity ratio ± SEM. of the indicated FRET pairs in fully occupied kinetochores. The number of kinetochores measured is indicated above the bars. All data are from N ≥ 3 experiments. Statistical significance was evaluated by a Mann Whitney test, ns = not significant; * = p<0.05; ** = p<0.01; *** = p<0.001; **** = p<0.0001

It is also noteworthy that the proximity ratio at the centromeric end of Ndc80C was only modestly reduced in unattached kinetochores. This observation suggests that the multivalent recruitment of Ndc80C by CenpT is mainly responsible for clustering at the centromeric ends of Ndc80C molecules (Huis In ’t Veld et al., 2016). Moreover, inter-complex FRET between Hec1-C and Spc25-C was also detectable in unattached kinetochores (Fig. 4B). Therefore, the longitudinal staggering of Ndc80C molecules likely results from the multivalence of CenpT.

To understand the influence of multivalent interaction of CenpT with Ndc80C in organizing the kinetochore, we adopted a comparative approach. In budding yeast, the centromeric protein linkages that recruit Ndc80C are monovalent, i.e. each recruits only one Ndc80C molecule (Dimitrova et al., 2016; Hornung et al., 2014; Malvezzi et al., 2013). Even though the CenpT homolog is present in budding yeast, it does not bind Ndc80C prior to anaphase (Bock et al., 2012; Dhatchinamoorthy et al., 2017; Malvezzi et al., 2013; Schleiffer et al., 2012). We expected that this key difference in the centromeric organization in human and yeast cells will give rise to distinct microtubule-dependent and independent aspects in their respective kinetochore architectures. Accordingly, in yeast kinetochores the centromeric ends of Ndc80C molecules were clustered during attachment, similar to human kinetochores, but this clustering vanished in unattached yeast kinetochores (compare the Spc25-C/Spc25-C measurements in human kinetochores with the Spc24-C/Spc24-C measurements in yeast; Fig. 4C, 4D & S7). Furthermore, in both attached and unattached yeast kinetochores Hec1-C/Spc25-C FRET was undetectable, which is consistent with a lack of Ndc80C staggering. The centromeric end of Mis12C (marked by N-Mis12 in the human kinetochore and N-Dsn1 in the yeast kinetochore; (Aravamudhan et al., 2014; Dimitrova et al., 2016; Petrovic et al., 2016)) showed a significant degree of clustering in budding yeast kinetochores, but not in human kinetochores. This clustering decreased in unattached budding yeast kinetochores, although the significance of this observation remains unknown. Interestingly, the degree of clustering of the microtubule-binding ends of Ndc80C molecules was similar in both kinetochores, implying that their organization on the microtubule lattice is similar. Furthermore, Ndc80C clustering was strongly dependent on attachment to the microtubule plus-end in both yeast and human kinetochores.

These data reveal the influence of different centromeric organizations on the protein architecture of human and budding yeast kinetochores. The differences between the two architectures strengthen our proposal that the multivalent CenpT linkage is the main source of centromeric clustering and longitudinal staggering of Ndc80C molecules in human kinetochores. Importantly, despite these differences both kinetochores achieve similar organization in the microtubule-binding ends of Ndc80C in the presence and absence of microtubule attachment.

### Centromeric tension and dynamic microtubules promote Ndc80C clustering

The sensitivity of Ndc80C clustering to microtubule attachment prompted us to study whether Ndc80C architecture in human kinetochores is also sensitive to centromeric tension. Centromeric tension arises from the opposing forces generated by bioriented sister kinetochores. To reveal the relationship between Ndc80C clustering and centromeric tension, we plotted inter-complex FRET between Ndc80C molecules at their microtubule binding ends (N-Nuf2/N-Nuf2) and at their centromeric ends (Spc25-C/Spc25-C) against the sister kinetochore separation, a proxy for the centromeric tension (referred to as the K-K distance, Fig. 5A). We found a positive linear correlation between FRET and K-K distance at both ends of Ndc80C (Fig. 5A). Thus, the proximity between Ndc80C molecules increases with centromeric tension at both the point of Ndc80C’s contact with the microtubule lattice and near its centromeric end.

**Figure 5.**
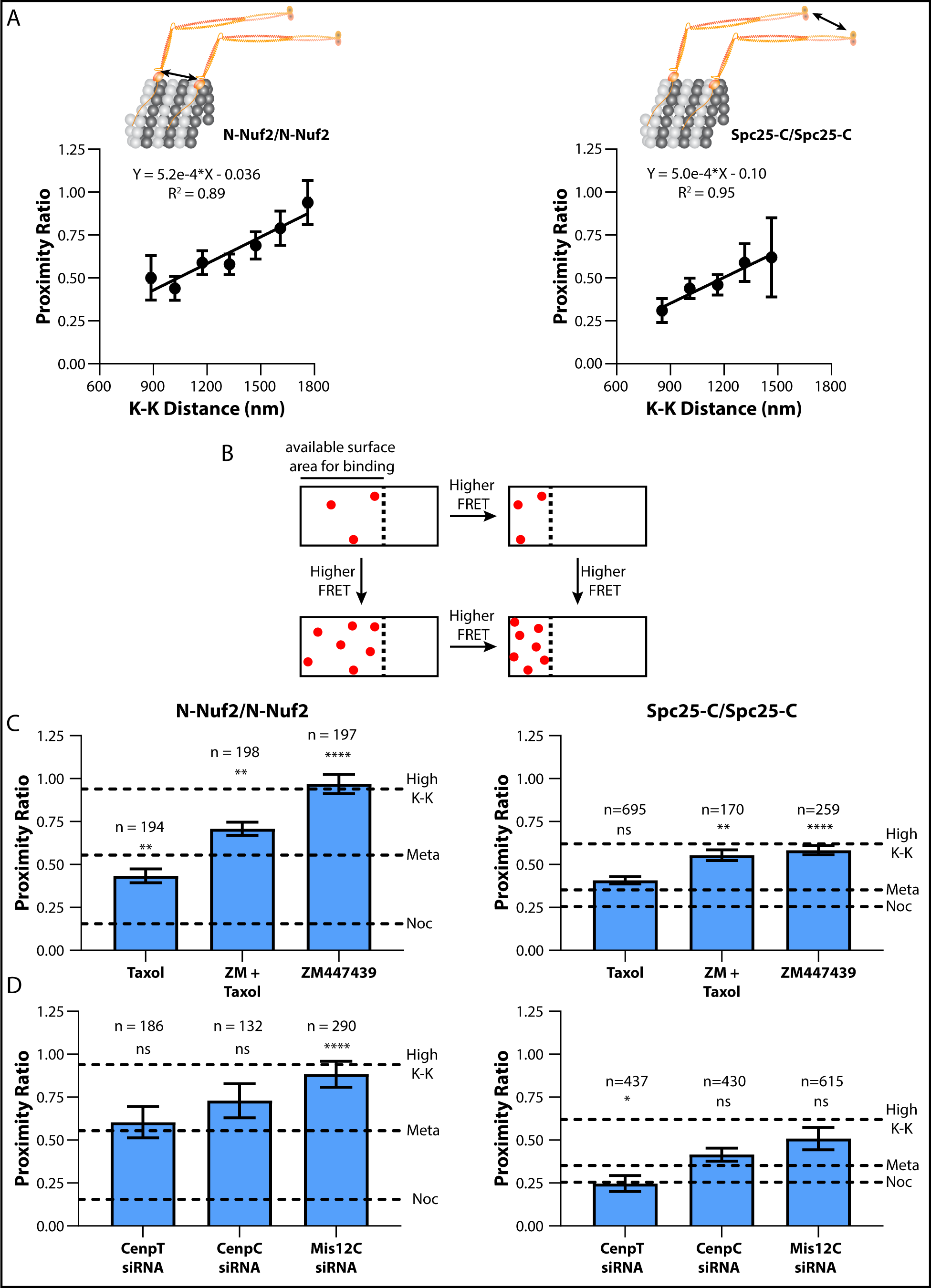
FRET between Ndc80C molecules correlates with centromeric tension. A) Proximity ratio measurements from fully occupied, metaphase kinetochores in either N-Nuf2/N-Nuf2 (left) or Spc25-C/Spc25-C (right) expressing HeLa cells were binned by their sister kinetochore separation distance, plotted, and fit with a linear regression. (Data are mean ± SEM; for N-Nuf2/N-Nuf2, n = 22, 49, 82, 77, 61, 28, 19; for Spc25-C/Spc25-C, n = 62, 76, 83, 32, 11) B) The diagram depicts the relationship between the binding density of Ndc80C and the FRET produced by a kinetochore. White rectangles represent the microtubule lattice, where the dashed line demarcates the region of the microtubule plus-end available for Ndc80C binding. Red dots represent bound molecules. C) N-Nuf2/N-Nuf2 and Spc25-C/Spc25-C FRET in response to the microtubule-stabilizing drug Taxol, the Aurora B kinase inhibitor ZM447439, or both. D) N-Nuf2/N-Nuf2 and Spc25-C/Spc25-C FRET after siRNA mediated knockdown of members of the Ndc80C recruitment pathways. In C) and D) data are the average proximity ratio ± SEM. The dashed lines indicate the average proximity ratio for untreated metaphase (Meta), nocodazole-treated (Noc), and for high tension kinetochores (High K-K). The number of kinetochores measured is indicated above the bars. All data are from N ≥ 3 experiments. Statistical significance between untreated, metaphase cells and each of the measurements in C) and D) was evaluated by a Mann Whitney test, ns = not significant; * = p<0.05; ** = p<0.01; *** = p<0.001; **** = p<0.0001

The clustering of Ndc80C molecules at their microtubule-binding end can increase in response to tension because: (1) the number of microtubule-bound Ndc80C molecules increases, (2) the spacing between bound molecules decreases, or both (Fig. 5B). Phosphoregulation of Ndc80C by the Aurora B kinase has been shown to increase the fraction of molecules bound to the microtubule lattice (Yoo et al., 2018). The average spacing between Ndc80C molecules can also change due to protein-protein interactions (e.g., oligomerization, accessory microtubule-binding proteins, etc.), the available binding surface on the microtubule, and the total number of microtubules per kinetochore (Alushin et al., 2012; Alushin et al., 2010; Helgeson et al., 2018; Janczyk et al., 2017; McEwen et al., 1997).

To understand the role of Aurora B mediated phosphoregulation on Ndc80C clustering, we treated HeLa cells with ZM447439, a small molecule inhibitor of the Aurora B kinase. ZM447439 treatment increased inter-complex FRET at both N-Nuf2 and Spc25-C such that the average value of the proximity ratio was equivalent to its value in kinetochores under the highest centromeric tension (Fig. 5C). ZM447439 treatment did not affect the range of K-K separation as compared to untreated cells, eliminating any potential role of tension on this experiment (Fig. S8; (Yoo et al., 2018)). Thus, an increase in the number of microtubule-bound molecules results in more clustering of Ndc80C molecules at the plus-end.

To reveal the role of microtubule dynamicity in clustering Ndc80C, we treated cells with Taxol, a microtubule stabilizing drug. Due to the dampening of tubulin polymerization dynamics by Taxol, the kinetochore-bound plus-ends become stabilized, and the number of bound microtubules increases (Fig. S9; (Fanara et al., 2004; Kumar, 1981; McEwen et al., 1997)). Therefore, Ndc80C molecules will have a larger microtubule surface area for binding. Accordingly, we measured a small decrease in inter-complex FRET at N-Nuf2 in Taxol-treated cells as compared to untreated metaphase cells (Fig. 5C and Fig. S9). Interestingly, FRET at Spc25-C did not change in the presence of Taxol, showing that microtubule stabilization has little effect on the organization of the centromere-anchored ends of Ndc80C molecules.

We also quantified inter-complex FRET in cells simultaneously treated with Taxol and ZM447439. The resulting FRET at N-Nuf2 was intermediate between what we measured with either Taxol or ZM447439 alone (Taxol = 0.43 ± 0.04 (S.E.M.), ZM447439 = 0.97 ± 0.06, Taxol + ZM447439 = 0.71 ± 0.04; Fig. 5C). Under this same condition the inter-complex FRET at Spc25-C was the same as that observed with ZM447439 treatment alone, consistent with the observation that Taxol does not influence the clustering of the centromeric ends of Ndc80C molecules. Overall, these observations show that the number of microtubule-bound Ndc80C molecules and, to a lesser extent, microtubule dynamics greatly influences the relationship between centromeric tension and Ndc80C clustering.

### Kinetochores depleted of Ndc80C recruitment linkages maintain Ndc80C clustering and form load-bearing microtubule attachments

Finally, we attempted to disrupt the Ndc80C clustering observed in metaphase kinetochores by knocking down either CenpT, CenpC, or Mis12C using siRNA. With these experiments, we wanted to ascertain the role of CenpT in clustering the centromeric ends of Ndc80C, and test whether the clustering of the microtubule-binding ends of Ndc80C is functionally significant. These knockdowns resulted in the expected ∼ 60-70% reduction in the amount of Ndc80C per kinetochore as previously reported (Fig. S8; (Suzuki et al., 2015)). In siRNA treated cells, several kinetochores remained unaligned, but most aligned at the metaphase plate and produced K-K distances similar to aligned kinetochores in untreated cells (Fig. S8). Only these aligned kinetochores were analyzed for fluorescence intensity and FRET quantitation.

We first examined the clustering of the centromeric ends of Ndc80C molecules by quantifying the inter-complex FRET between Spc25-C/Spc25-C. These measurements indicated that the clustering of the centromeric ends of Ndc80C was not significantly influenced by CenpC and Mis12C siRNA (Fig. 5D). However, we saw a modest decrease in FRET upon CenpT depletion. This is consistent with the recruitment of up to 3 Ndc80C molecules per CenpT molecule (Huis In ’t Veld et al., 2016). We next assessed the clustering of the microtubule-binding ends of Ndc80C molecules by quantifying the inter-complex FRET between N-Nuf2/N-Nuf2. We expected that the reduced number of Ndc80C molecules per kinetochore will reduce its surface density on the centromere, and hence its ability to cluster. Surprisingly, we found that rather than decreasing, inter-complex FRET between N-Nuf2/N-Nuf2 either remained unchanged or increased significantly (Fig. 5D, also see Fig. S8A). In the case of Mis12C knock-down, the inter-complex FRET approached the value measured during ZM447439 treatment. As the ZM447439 measurement likely represents maximal microtubule binding of Ndc80C molecules, these data suggest that a greater fraction of Ndc80C molecules actively bind to microtubules under the siRNA condition. Thus, despite significant reductions in Ndc80C copy number per kinetochore, the remaining pool of Ndc80C molecules effectively bound microtubules to form load-bearing attachments.

## Discussion

Our analysis adds a crucial new dimension to the emerging model of human kinetochore architecture by defining the distribution of key protein molecules around the plus-end and along the longitudinal axis of an attached microtubule. We synthesized this information with protein structures and interactions to construct a detailed model of the organization of human kinetochore-microtubule attachments (Fig. 6A). In synthesizing this model, we considered the structure and interactions of the human kinetochore’s repeating ∼ 26-subunit core (Pesenti et al., 2018; Weir et al., 2016). This core is seeded by centromeric nucleosomes containing the centromere-specific histone CenpA, which anchor the CenpT and CenpC recruitment linkages for Ndc80C (Gascoigne et al., 2011; Klare et al., 2015; McKinley et al., 2015; Pesenti et al., 2018; Weir et al., 2016). Therefore, the number and spatial distribution of CenpA nucleosomes will play a critical role in the centromeric distribution of CenpC and CenpT molecules. However, due to technical challenges, we did not analyze either CenpA or CenpC in this study. Nonetheless, current estimates suggest that at least 44 CenpA nucleosomes per centromere participate directly in nucleating the human kinetochore (Bodor et al., 2014; Logsdon et al., 2015). Because one CenpA nucleosome recruits two copies of the CCAN, this number suggests that the human kinetochore should contain ∼ 80 copies of each CCAN subunit (Weir et al., 2016). Our quantitation of CenpHIK is consistent with this prediction (Fig. 3D). Since the number of microtubules per kinetochore is ∼ 17-20, we depict two CenpA nucleosomes contributing to the attachment of a single microtubule in our model (Fig. 6A). We find that the protein linkages nucleated by the CenpA nucleosomes are sparsely distributed within the kinetochore with a minimum distance of ∼ 10 nm between them. Despite this sparse distribution, the multivalence of CenpT ensures that a significant fraction of Ndc80C molecules are clustered at their centromeric ends. This CenpT-mediated clustering of Ndc80C and the many flexibilities in the linkage leading up to Ndc80C are likely to promote their ability to persistently attach to the dynamic plus-end (Volkov et al., 2018).

**Figure 6.**
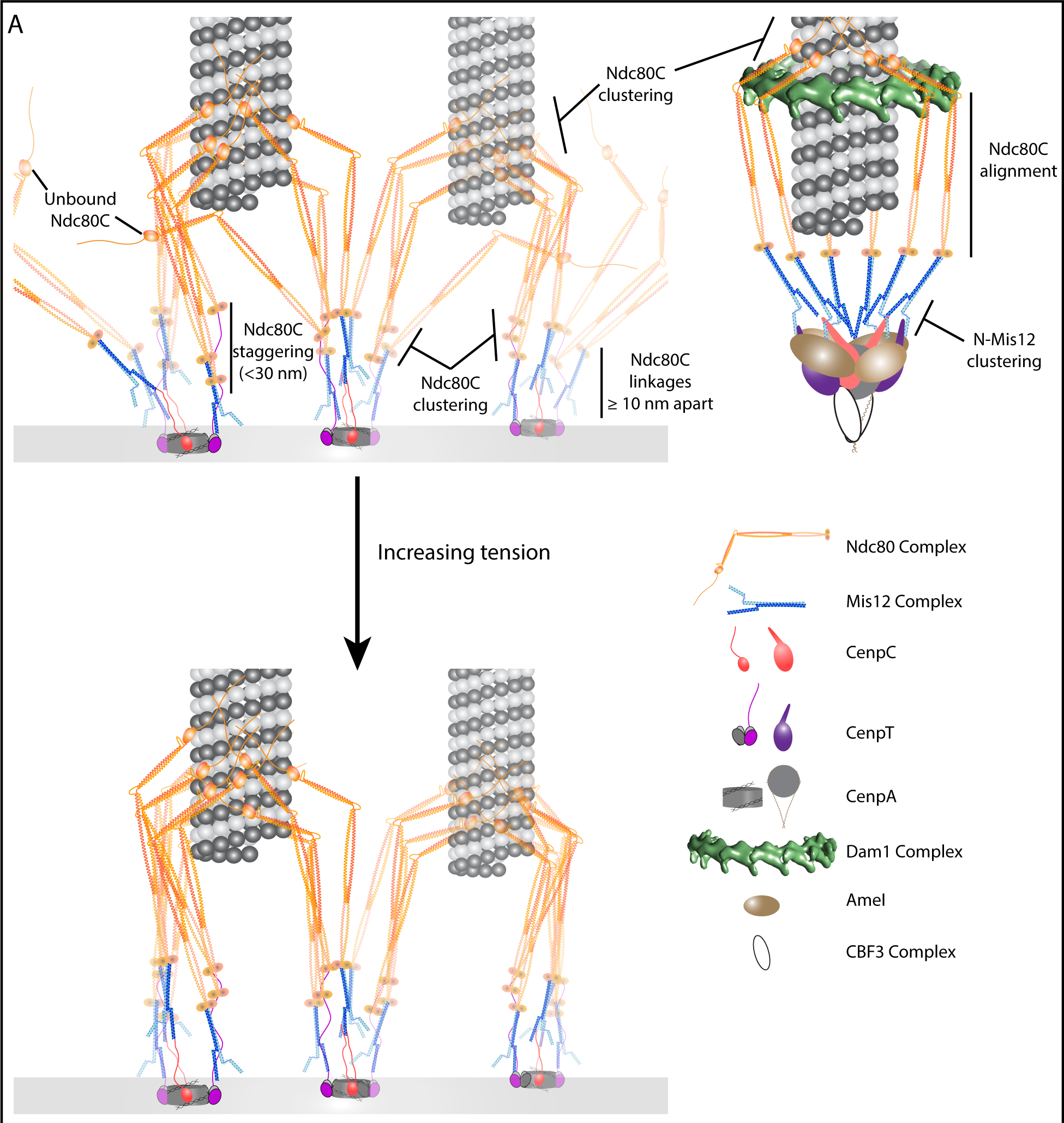
Architectural models of human and budding yeast kinetochore microtubule attachments. The protein organization of human kinetochore-microtubule attachment sites (left) is responsive to physical attachment to the microtubule lattice and to centromeric tension, both of which act to increase the density of microtubule-bound Ndc80C molecules. For comparison, we include a model of the budding yeast kinetochore (upper right). The legend (bottom right) identifies proteins for both models. Key architectural details are emphasized. See text for further details.

The sensitivity Ndc80C FRET to microtubule attachment reveals the adaptability of kinetochore architecture to its mechanical state. Upon attachment, the microtubule-binding domains of Ndc80C molecules transition from a state of little-to-no clustering to a state of high clustering (Fig. 2 and Fig. 4). Furthermore, this clustering increases proportionally with centromeric tension (Fig. 5). These data highlight the importance of the microtubule lattice for scaffolding the clustered organization of Ndc80C molecules and suggest that centromeric tension may be correlated with the number of microtubule-bound Ndc80C molecules. This hypothesis is supported by the significant increase in Ndc80C clustering upon Aurora B kinase inhibition, which promotes maximal binding of Ndc80C molecules at the kinetochore (Fig. 5C; (Yoo et al., 2018)). It should be noted that Aurora B inhibition may also affect the function of key microtubule-binding proteins (e.g. the Astrin-SKAP complex) that interact with Ndc80C, and thereby indirectly influence the distribution of Ndc80C (Manning et al., 2010; Redli et al., 2016; Schmidt et al., 2010). Microtubule dynamics also play a role in Ndc80C clustering. This is most clearly seen when Aurora B activity and microtubule dynamics are inhibited simultaneously: Ndc80C clustering decreases at its microtubule-binding domains compared to Aurora B inhibition alone (Fig. 5C). While the precise mechanism by which microtubule dynamics affect Ndc80C clustering is unclear, it is likely that the dynamicity limits Ndc80C distribution to a smaller region near the plus-end (Long et al., 2017).

Given the influence of microtubule binding on Ndc80C clustering, we were surprised that clustering either remained unaffected or increased upon siRNA-mediated depletion of the centromeric receptors of Ndc80C (Fig. 5D). Furthermore, these depleted kinetochores were capable of forming load bearing attachments (Fig. S8 &S9). These data suggest that within the depleted kinetochores a higher fraction of the remaining Ndc80C molecules participated in engaging the plus-end. Because of the smaller number of Ndc80C molecules per kinetochore, it is possible that the average force experienced by each molecule is higher than the force per molecule in normal metaphase kinetochores (Fig. S8A). The higher tension per molecule will in turn give rise to higher FRET. Pleiotropic changes in kinetochore regulation will also affect these measurements (Caldas et al., 2013). Nonetheless, these findings suggest that multiple CenpA nucleosomes may contribute Ndc80C molecules to engage with the plus-end of a single microtubule (Fig. 6A). Thus, even though the Ndc80C molecules in the human kinetochore are recruited by repeating subunits nucleated by individual CenpA nucleosomes, they operate as a ‘lawn’ of microtubule-binding domains (Dong et al., 2007; Zaytsev et al., 2015; Zaytsev et al., 2014).

Finally, our study highlights the similarities and differences in the architectures of kinetochores built upon point centromeres in budding yeast and regional centromeres in HeLa cells. Unlike the human kinetochore, the yeast kinetochore is nucleated by just one CenpA nucleosome, and it forms a persistent attachment with only one microtubule (Fig. 6A; (McIntosh et al., 2013)). Therefore, all Ndc80C molecules in a budding yeast kinetochore interact with the same microtubule plus-end. Consistent with this picture, both the microtubule-binding domains of Ndc80C molecules and the centromere-binding ends of the Mis12C molecules cluster together in the yeast kinetochore (Fig. 4D; (Aravamudhan et al., 2014; Dimitrova et al., 2016)). In contrast, only Ndc80C molecules cluster in the human kinetochore; Mis12C molecules do not (Fig. 4B). Furthermore, adjacent Ndc80C molecules are aligned with one another in the yeast kinetochore, but they stagger along the longitudinal axis of the human kinetochore. In yeast kinetochores, the absence of Ndc80C staggering is likely enforced by the point centromere as well as the Dam1 ring or ring-like structure (Ng et al., 2019). In human kinetochores, the staggered organization of Ndc80C arises from the multivalence of CenpT and the possibility of Ndc80C linkages anchored to different CenpA subunits contacting the same plus-end. We estimate that the staggering between adjacent Ndc80C molecules is no greater than 30 nm, although we cannot rule out the possibility that non-adjacent Ndc80C molecules are staggered by even larger distances. The staggered arrangement of Ndc80C molecules will enhance the persistence of human kinetochore-microtubule attachments and their ability to track the tips of microtubule plus-ends (Hill, 1985). Given the significant differences in the organizations of human and yeast centromeres, it is remarkable that both kinetochores achieve a similar degree of clustering at the microtubule-binding ends of their Ndc80C molecules. Thus, it is possible that this similarity in the kinetochore-microtubule interfaces of yeast and humans represents a generally conserved feature of kinetochore architecture.

## Acknowledgements

The authors wish to thank the Single Molecule Analysis in Real-Time (SMART) Center of the University of Michigan, seeded by NSF MRI-R2-ID award DBI-0959823 to Nils G. Walter, as well as Damon Hoff for training, technical advice and use of the ALBA time-resolved confocal microscope.

## Author Contributions

Conceptualization, Methodology, Writing – Original Draft, Writing – Review & Editing, Visualization, and Supervision, A.A.K. and A.J.P.; Formal Analysis and Investigation, A.A.K., S.K., and A.J.P.; Software, Resources, and Funding Acquisition, A.J.P.

## STAR Methods

### Construction of stable protein expression HeLa cells

For the co-expression of fluorophore-tagged kinetochore proteins, HeLa cell lines were generated containing a stable chromosomal insertion of a dual-expression vector. The HeLa A12 cell line (gift from the Lampson lab) contains a lentiviral-based chromosomal insertion of a pair of incompatible Cre/Lox sites in front of the human EF-1α promoter (see (Khandelia et al., 2011) for details). Using standard molecular cloning, we created cassettes capable of Cre recombinase-mediated integration at this chromosomal locus that were based on the pERB131 plasmid backbone (gift from the Lampson lab). Briefly, the pERB131 backbone contains two open-reading frames (ORFs), one that becomes under the control of the constitutive EF-1α promoter upon successful integration (ORF1) and a second (ORF2) which is controlled by a tetracycline responsive promoter (Tet-ON). All proteins examined in this study were cloned into one of these two ORFs. The cassette also contains a gene for puromycin resistance which aided in the selection of HeLa cells with successful integration. All HeLa cell lines generated for this study are listed in Supplementary Table 2.

Integration was performed using the Lipofectamine 3000 Reagent kit (Thermo Fisher Scientific) to co-transfect cells with the pERB131 cassette of interest and a Cre-expression plasmid ((gift from the Lampson lab). Two days post-transfection, 2 µg/mL puromycin was added to the cell media for selection over the course of two weeks. Successful transformants were then maintained in media containing 1 µg/mL puromycin.

### Fluorescence Microscopy

All fluorescence and FRET imaging was performed on a Nikon Ti-E inverted microscope with a 1.4 NA, 100x, oil immersion objective. A Lumencor LED light engine (472/20 nm GFP excitation, 543/20 nm mCherry excitation) served as the laser power source. All filters are from Chroma and included: 1) a dual-band excitation filter ET/GFP-mCherry (59002x); 2) an excitation dichroic (89019bs); 3) an emission-side dichroic (T560lpxr); 4) and emission filters ET525/50m and ET595/50m. Images were acquired on an Andor iXon3 EMCCD camera (pixel size = 160 nm). Cell images were either 20 or 10 plane z-stack image series for HeLa and budding yeast cells, respectively. The step size between planes was 0.25 µm. For most experiments, the acquisition rate for GFP and mCherry was set at 400 ms. Occasionally, when the copy number of fluorophore-tagged proteins was low (e.g., CCAN proteins or during siRNA mediated knockdowns) the acquisition rate was increased to obtain higher fluorescence signal. A simple linear correction was applied to normalize fluorescence intensity values to a 400 ms acquisition rate. To account for fluctuations in laser power or other artifacts in our microscopy setup, we collected images of ∼ 20 anaphase budding yeast cells expressing Ndc80-GFP and Spc25-mCherry before all experiments. Since budding yeast incorporate a stable number of proteins per kinetochore, any changes in GFP and mCherry brightness should be a result of instrument-derived fluctuations. In this way, ratiometric correction factors were derived for each day of imaging to normalize all FRET measurements throughout the course of this study.

For HeLa, cells were plated in multi-chamber glass-bottomed dishes (Lab-Tek®II) in DMEM media (Gibco) supplemented with 10% FBS (Corning), and 100 U/mL penicillin and 100 µg/mL streptomycin (Gibco). Cells were treated with 1 – 2 µg/mL doxycycline for 48 hrs to induce the expression of ORF2 proteins. Treatments with siRNA were performed using the Lipofectamine RNAiMAX kit (Invitrogen), using 30 pmol of each protein-specific siRNA and an incubation period of at least 48 hrs (siRNAs listed in Supplementary Table 3). During imaging, cell media was changed to DMEM without any phenol red supplemented with 10% FBS and 100 U/mL penicillin and 100 µg/mL streptomycin. During imaging, the microscope stage is fitted with a heated chamber with CO_2_ respirator and objective warmer (Live Cell Instrument). For several experiments, we employed double-thymidine synchronization with 2.5 mM thymidine. For imaging unattached kinetochores, cells were treated with 100 ng/mL nocodazole and incubated at least 30 min before imaging. For imaging attached, tensionless kinetochores, cells were treated with 10 µM Taxol and incubated at least 10 min before imaging. For experiments with the Aurora B inhibitor, ZM447439, we added 10 µM MG132 and incubated 5 min before adding 3 µM of the ZM447439 drug. Cells were incubated an additional 10 min before imaging. For imaging with both ZM447429 and Taxol, the same procedure was followed as above, adding MG132 and ZM447439 first, incubating 10 min, then adding Taxol. Attached kinetochores were distinguished from unattached kinetochores by their positioning at the spindle mid-zone. Unattached kinetochores were often dispersed through the cell body with random orientations with respect to the spindle and greatly reduced (∼ 800 nm) sister kinetochore separation.

For budding yeast, cells were grown at 30 °C to mid-log phase in yeast peptone (YP) media supplemented with 2% glucose. For strains with galactose-inducible promoters, the YP media was supplemented with 2% raffinose and varying concentrations of galactose. The appropriate galactose concentration was determined as that which produced average fluorescence signals at kinetochores that were equal to the fluorescence signal in strains without inducible-promoters. Prior to imaging, cells were rinsed and concentrated in synthetic drop-out media. For imaging of unattached kinetochores, mid-log phase cells were treated with 15 µg/mL nocodazole for 1.5 hrs before rinsing and concentrating cells in synthetic media supplemented with 15 µg/mL nocodazole. Metaphase kinetochore clusters were designated by sister pairs with a separation of ∼ 0.8 to 1 µm. For nocodazole-treated cells, unattached kinetochores were identified as the dimmer fluorescent puncta separate from the brighter, spindle-localized attached kinetochores. Cells were imaged on 22x22 mm glass coverslips. All yeast strains used in this study are from (Aravamudhan et al., 2014).

### FRET Quantification and Image Analysis

To measure FRET, a semi-automated graphical user interface written in MATLAB was used to analyze cell images. The implementation of this program is described in (Joglekar et al., 2013). The raw FRET intensity, measured as the fluorescence intensity observed in the mCherry channel upon excitation with the GFP-specific laser, contains contaminating signal from GFP bleed-through and mCherry cross-excitation. The contribution of these signals was measured in HeLa cells expressing either Spc25-GFP or Spc25-mCherry alone (Fig. S3; GFP bleed-through = 5.79 ± 0.17%, mCherry cross-excitation = 6.64 ± 0.18%). Subtracting these values from the raw FRET intensity yields the sensitized emission due to FRET. Given the dynamicity of the kinetochore and differing stoichiometries between its protein subunits, the sensitized emission was further normalized by the sum of the GFP bleed-through and mCherry cross-excitation. Since the bleed-through and cross-excitation are proportional to the number of fluorophore-tagged molecules, this normalization essentially yields the sensitized emission/molecule, a metric we refer to as the proximity ratio:

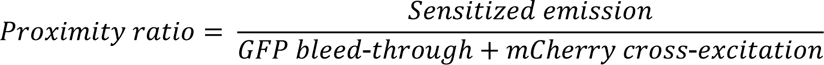

As noted in the main text, we observed considerable variation in the fluorophore intensity of both HeLa and budding yeast kinetochores. The sources of these variations are likely due to multiple effects, e.g. incomplete fluorophore maturation, inefficient siRNA knockdowns, and variations in fluorophore intensity with depth below the coverslip. Different approaches were taken in HeLa and budding yeast cells to correct for these variations.

For HeLa cells, we measured the co-variation in GFP and mCherry signals at kinetochores in siRNA-treated cells co-expressing Spc25-GFP and Spc25-mCherry (Fig. 1C). A plot of the Spc25-mCherry v. Spc25-GFP fluorescence intensities measured at each kinetochore was generated after binning the values by the stoichiometric ratio of mCh:GFP. A linear regression fit to the plot yielded the equation y = -0.4418x + 11010. As Spc25 is a subunit of Ndc80C, we referred to (Suzuki et al., 2015) for the number of Ndc80C molecules per HeLa kinetochore (∼ 244). Using this information and the maximal GFP and mCherry values determined by the linear regression fit, we determine the single molecule brightness of GFP and mCherry to be 102.1 ± 5.0 and 45.1 ± 0.9 a.u. The single molecule brightness of GFP and mCherry were used to determine to the number of molecules at single kinetochores. These values were filtered according to the protein counts and the error bounds from (Suzuki et al., 2015). For members of the CenpHIKM complex, we first noted that from unfiltered intensity values, the cumulative average of CenpH, CenpI, and CenpK was ∼ 86.2 molecules/kinetochore. Since the CenpHIKM complex aids in the recruitment of CenpT, the lowest copy number protein measured in this study, we set the lower limit of CenpHIKM molecules the same as for CenpT (i.e., 64 molecules at minimum per kinetochore). The upper limit was then set at 110 molecules/kinetochore to place the average value as the midpoint of these extremes (i.e., we used a range of 64-110 CenpHIKM molecules to filter for saturation of kinetochore binding). We note that for experiments involving siRNA-mediated knockdowns, the expected number of molecules is documented in (Suzuki et al., 2015). For nocodazole measurements, we adhered to the lower limits determined from (Suzuki et al., 2015), but did not place an upper limit on these values for two reasons: 1) it has been previously noted elsewhere and in our studies that nocodazole-treated, unattached kinetochores maintain greater numbers of molecules than attached, metaphase kinetochores; and 2) the disorganized spindles and lack of tension between sister kinetochores made it difficult to measure single kinetochores. Similarly, due to the lack of kinetochore tension upon taxol treatment, we filtered taxol measurements between 212-488 molecules. Therefore, these measurements may represent more than one kinetochore, although for the purposes of quantifying FRET this is not a problem due to the normalization inherent in the proximity ratio that we report.

For budding yeast, filtering metaphase kinetochores by absolute brightness was complicated by the fact that nocodazole treatment creates unattached kinetochore clusters of variable size. Therefore, for consistency between metaphase attached and the unattached nocodazole-treated kinetochores, all measurements were filtered to contain only data points with an mCh:GFP ratio of 0.5 – 2. All statistical analyses in this study were performed using the Mann-Whitney U test.

### Fluorescence lifetime imaging (FLIM) microscopy and determination of GFP lifetimes

FLIM data were collected on an ISS ALBA time-resolved laser-scanning confocal system. This setup consists of: 1) an Olympus IX-81 microscope with a U-Plan S-APO 60X 1.2 NA water immerision objective; 2) an SPC-830 time-correlated single photon counting (TCSPC) board (Becker & Hickl); 3) an SC-400-6-PP supercontinuum laser (Fianium); 4) and two photomuliplier tubes (PMT) detectors (Hamamatsu H7422P-40). During data collection, the objective was also equipped with a 37 °C temperature-controlled sleeve. HeLa cells were plated in 35 mm glass-bottomed dishes (MatTek) and imaged in DMEM media without any phenol red supplemented with 10% FBS and 100 U/mL penicillin and 100 µg/mL streptomycin. GFP and mCherry excitation were performed with 488 nm and 561 nm lasers, interleaved with 20 MHz frequency and 256 ADC resolution. The pixel-dwell time was 0.2 ms and laser power was adjusted to keep photon counts between 500,000 – 1,000,000 per pixel.

FLIM data were analyzed using the VistaVision software analysis program (ISS). To distinguish between cytosolic versus kinetochore-localized GFP, we employed intensity threshold masks. This method proved effective since kinetochore-localized GFP always provided higher counts/pixel than cytosolic GFP. Additionally, kinetochore pixels were further isolated by cropping the images to contain only the cellular region corresponding to the metaphase plate. At minimum, cytosolic GFP had a lower threshold of 30 photon counts/pixel to distinguish from background. We also maintained a buffer of at least 25 photon counts/pixel between the upper threshold for the cytosolic GFP and the lower threshold for the kinetochore-localized GFP to prevent cross-contamination of signals. After appropriate thresholding, photon counts from all pixels were summed. As the GFP excitation laser was pulsed first during the interleaved excitation, only photons collected between the first 6.6 – 24.8 ns of each pulse were included. GFP lifetimes were estimated by fitting the histograms of the photon arrival times to single-component exponential decays, using a software generated instrument response function (IRF). The FRET efficiency was calculated by comparing the difference between the GFP lifetime in the absence and presence of an mCherry acceptor 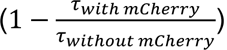. We note that the GFP lifetime in the absence of an mCherry acceptor was highly dependent on the protein subunit to which it was attached and on temperature. Therefore, for all FRET pairs the GFP lifetimes without an mCherry acceptor were measured independently.

### Western blot analysis

HeLa cell lysates were collected from cultures grown in 6-well plates (Corning), seeding at a density of ∼ 100K cells/well. Cells were grown for 3 days with the appropriate drug and siRNA treatments applied, after which cells were rinsed and aspirated. Lysates were collected in 50 µL SDS-PAGE buffer containing 2-mercaptoethanol using a cell scarper. Lysates were sonicated, denatured at 95 °C, vortexed and centrifuged before loading onto a 4% stacking/10% resolving SDS-PAGE gel. After running, PAGE gels were transferred to PVDF membranes (pre-activated by soaking in methanol) via electrophoresis in transfer buffer (1.4% glycine, 0.3% Tris-base in H_2_O). Blots were blocked with 5% milk in tris-buffered saline (TBS) for 30 min. and then incubated with primary antibody for 1 hr (primary antibodies prepared in 3% BSA + 0.1% Triton X100 + 01% NaN_3_ in phosphate-buffered saline). After rinsing, blots were incubated with secondary antibody prepared in 5% milk in TBS for 30 min. After rinsing, blots were developed via chemiluminescence (Immobilon Western reagent from Millipore) and exposure to x-ray film. Blot images were analyzed in FIJI. All antibodies used in this study are provided in Supplementary Table 4.

## Supplementary Figure Legends

**Figure S1.**
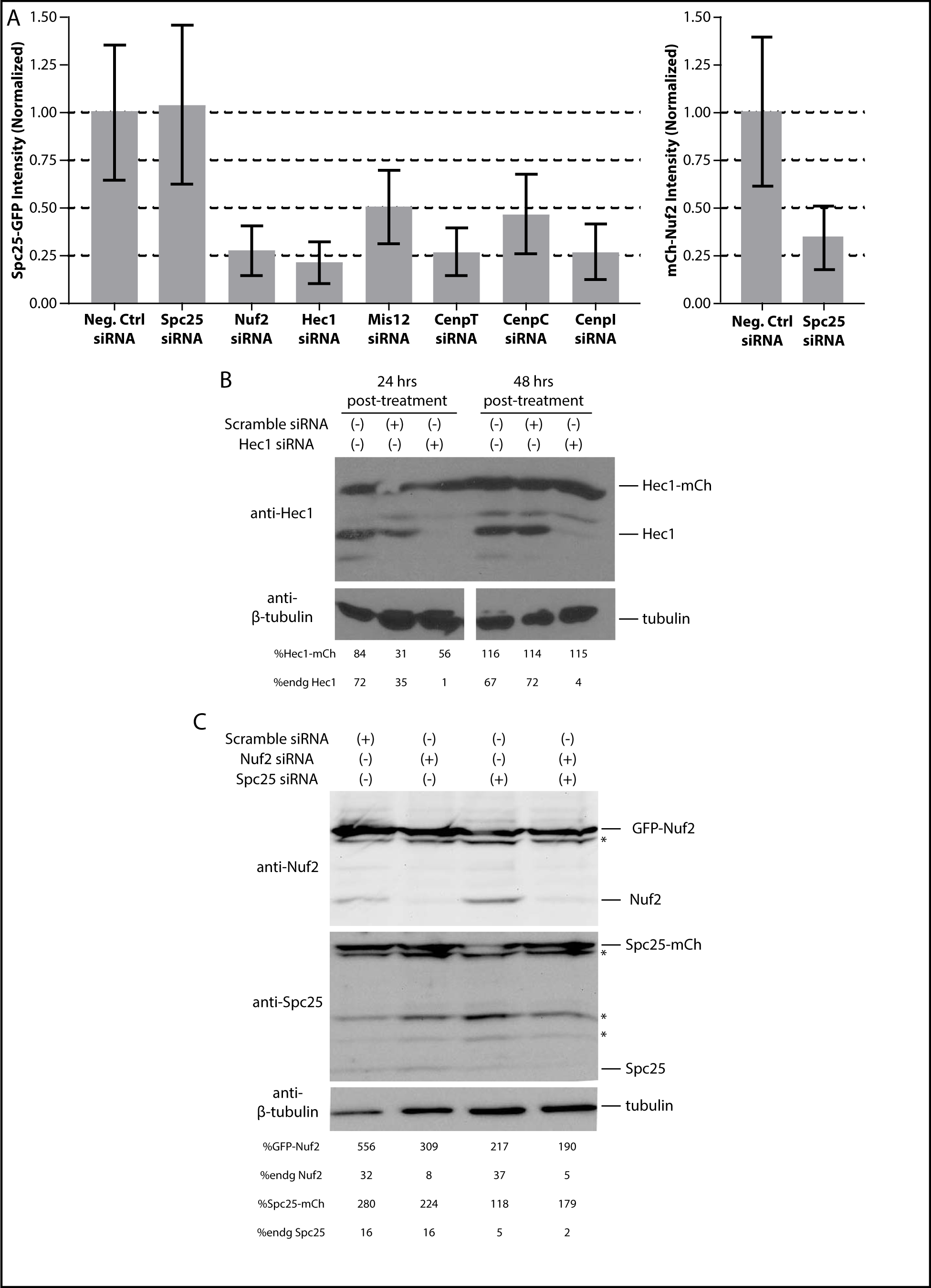
Knockdown of endogenous, unlabeled kinetochore proteins via siRNA treatment. A) The recruitment of siRNA resistant Spc25-GFP (left) or mCherry-Nuf2 (right) to kinetochores was affected by siRNAs targeting the indicated subunits which are either directly or indirectly involved in Ndc80C recruitment. Bars are the average fluorescence intensity ± S.D. All measurements are normalized to negative control siRNA treated cells. B) Western blot analysis of HeLa cells expressing siRNA resistant Hec1-mCherry. C) Western blot analysis of HeLa cells co-expressing siRNA resistant GFP-Nuf2 and Spc25-mCherry. Cell lysates were collected 48 hours post siRNA treatment. Numbers below the blots in B) and C) are %band intensities relative to the β-tubulin loading control for either the siRNA resistant, fluorophore tagged proteins or for the endogenous, unlabeled proteins. Asterisks indicate contaminating bands.

**Figure S2.**
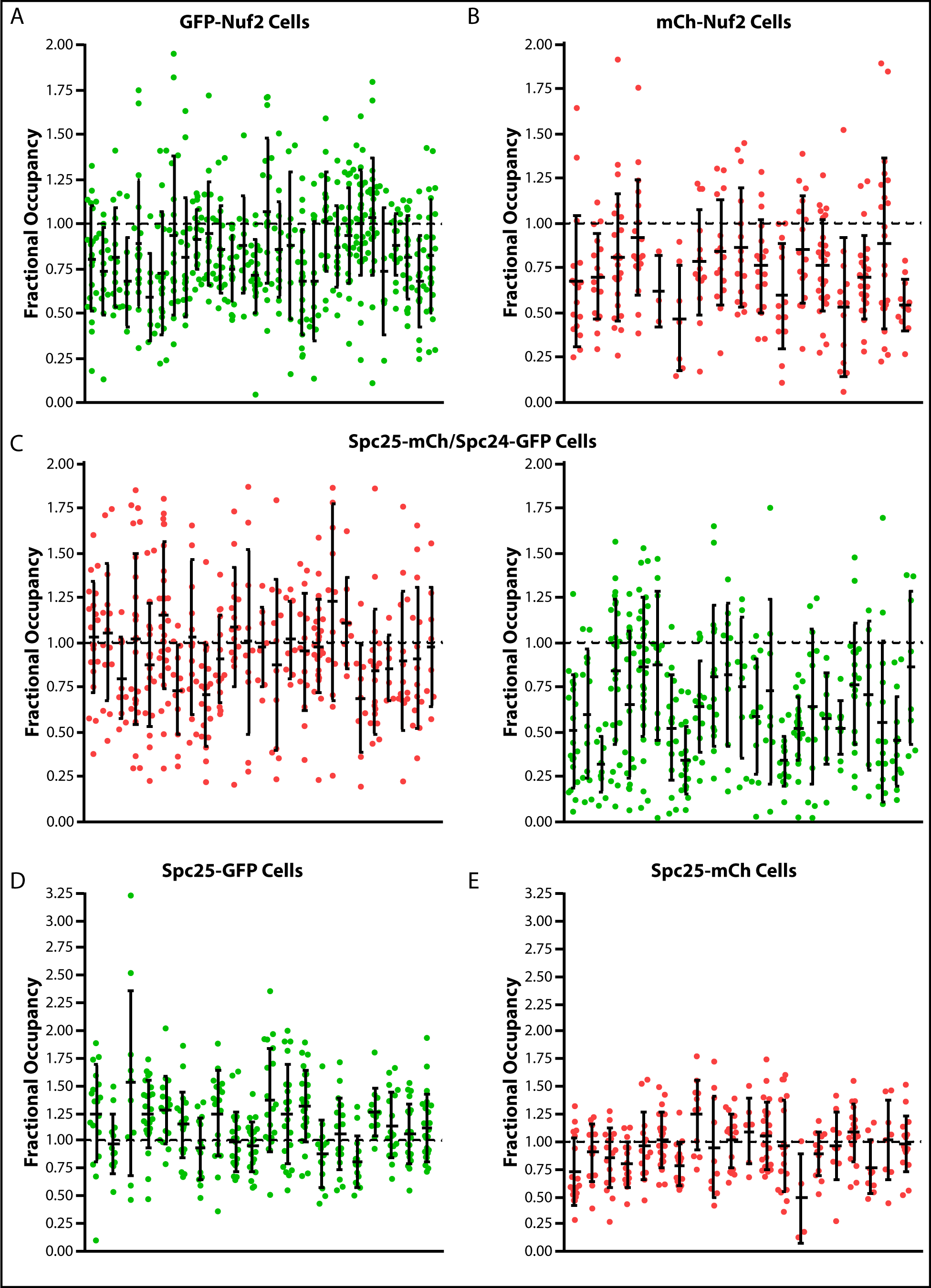
Variation in fluorophore intensities between individual kinetochores in HeLa cells. A) – E) Plots depict kinetochore fluorophore intensities normalized to the intensity value for 100% kinetochore occupancy. Groups of measurements come from single cells, and each point represents a single kinetochore. Error bars are the average kinetochore intensity for a single cell ± S.D. All cells were treated with siRNAs targeting the endogenous, unlabeled counterpart to the fluorophore tagged proteins. Note that in C), measurements are from HeLa cells co-expressing Spc25-mCherry and Spc24-GFP. All other plots are from HeLa cells expressing the indicated protein in isolation.

**Figure S3.**
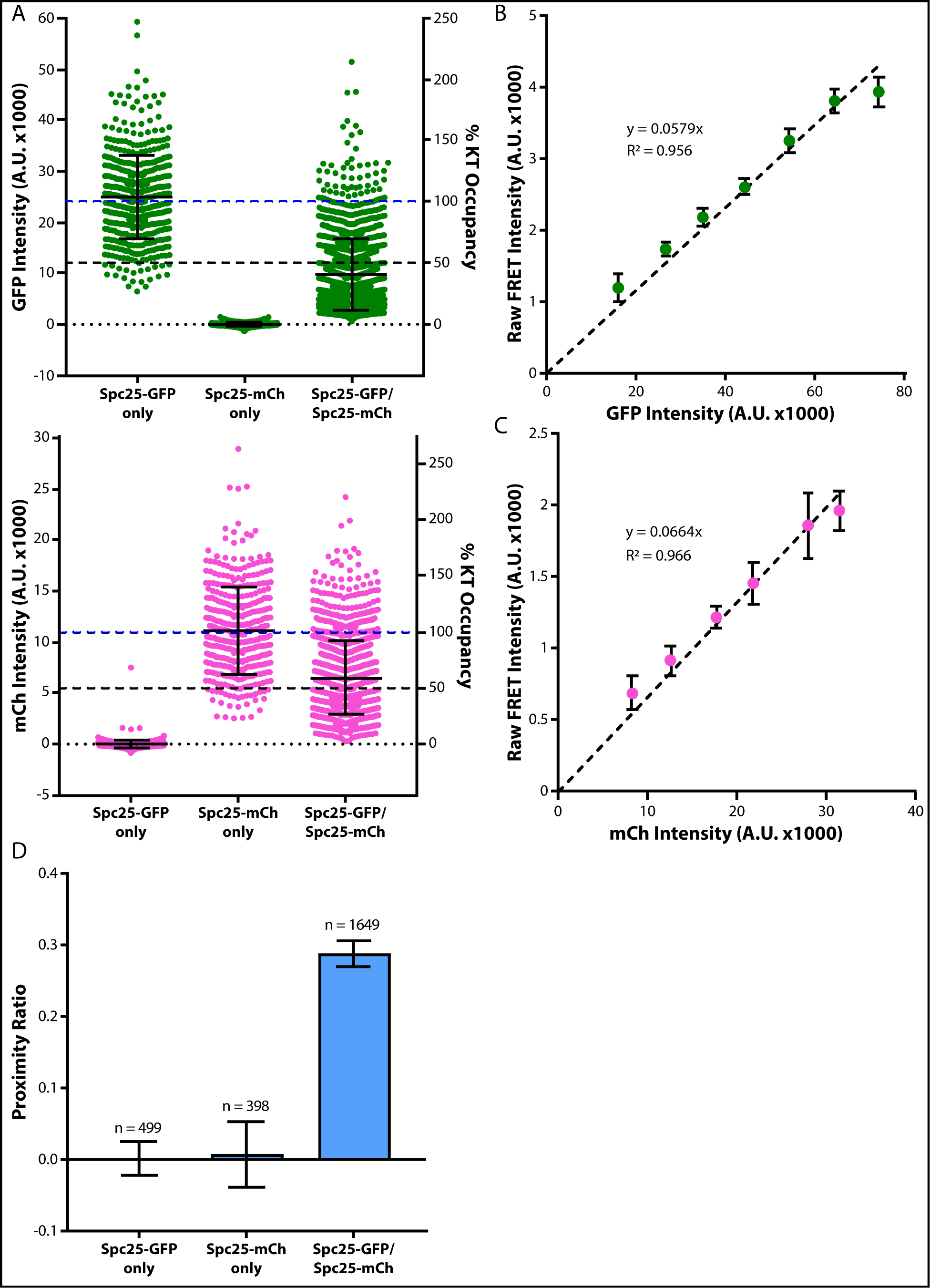
Calibrations for the measurement of FRET in live HeLa cells. A) GFP and mCherry intensities in HeLa cells expressing either Spc25-GFP, Spc25-mCherry, or co-expressing Spc25-GFP & Spc25-mCherry, after siRNA-mediated knockdown of the endogenous, unlabeled Spc25. Measurements are from unfiltered metaphase kinetochores. Data are average fluorescence intensity ± S.D. B) Donor emission bleed-through was estimated from the slope of a linear regression fit to a plot of raw FRET intensity versus GFP intensity in HeLa cells expressing Spc25-GFP in isolation. Data are binned by the GFP intensity. Error bars are ± S.E.M. C) Similar to B), but estimating acceptor cross-excitation from measurements in HeLa cells expressing Spc25-mCherry in isolation. D) The donor bleed-through and acceptor cross-excitation ensure FRET is only measurable when the donor and acceptor fluorophores are co-expressed. Bars are the average proximity ratio at metaphase kinetochores ± S.E.M. The number of measurements is indicated above the bars. Measurements are from kinetochores before filtering for 100% occupancy.

**Figure S4.**
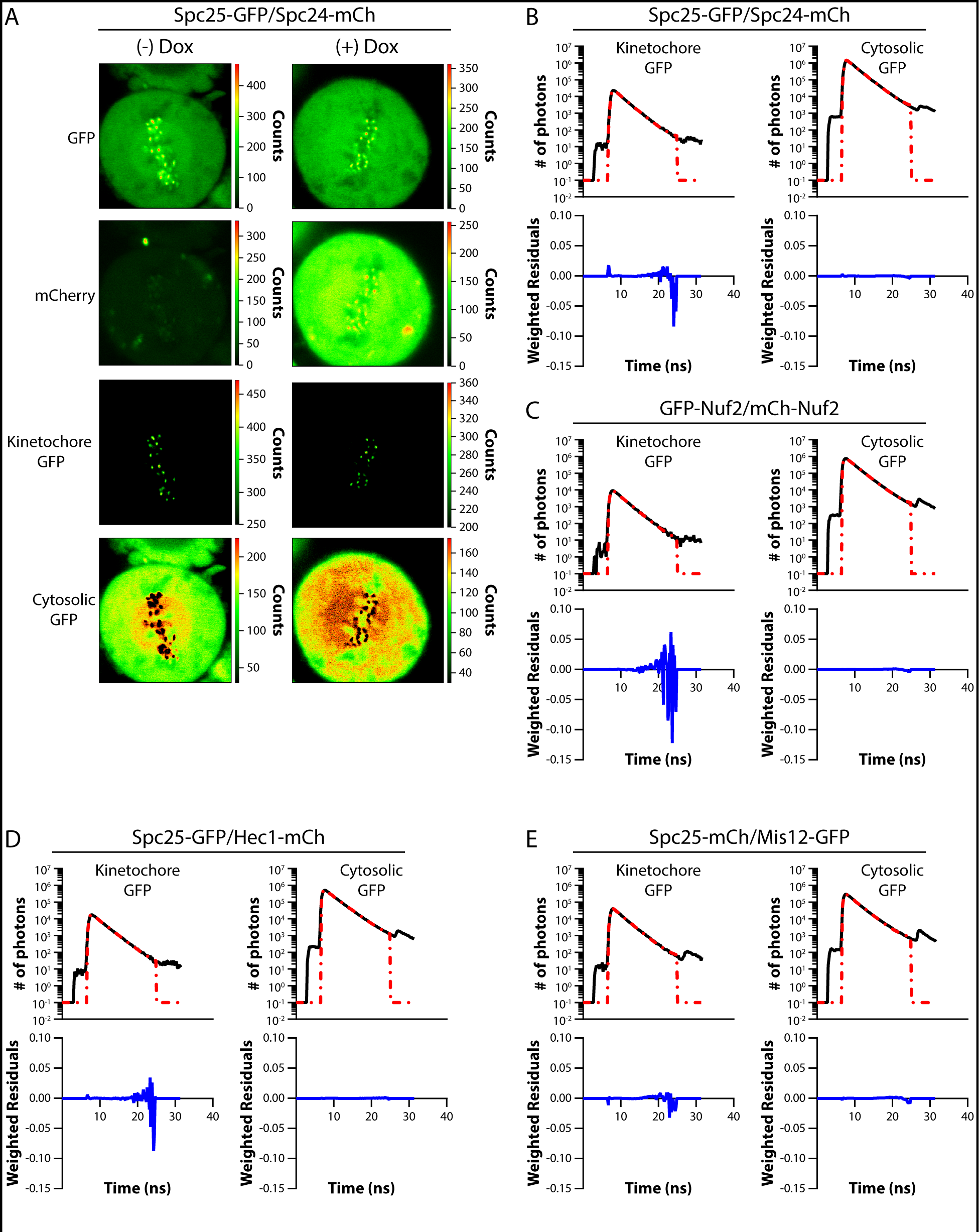
FRET-induced changes in donor lifetime measured by FLIM microscopy. A) Example confocal micrographs of Spc25-GFP/Spc24-mCherry HeLa cells for FLIM microscopy. Doxycycline (Dox) induces expression of the Spc24-mCherry protein. All images are scaled by the number of photons/pixel (scale to the right of images). Intensity thresholding was used to separate kinetochore-localized from cytosolic GFP pixels (bottom two rows). B) – E) Example photon arrival histograms for kinetochore-localized and cytosolic GFP in HeLa cells co-expressing the indicated proteins. Histograms are fit with single component exponential decays (red dashed lines). Weighted residuals are below histograms.

**Figure S5.**
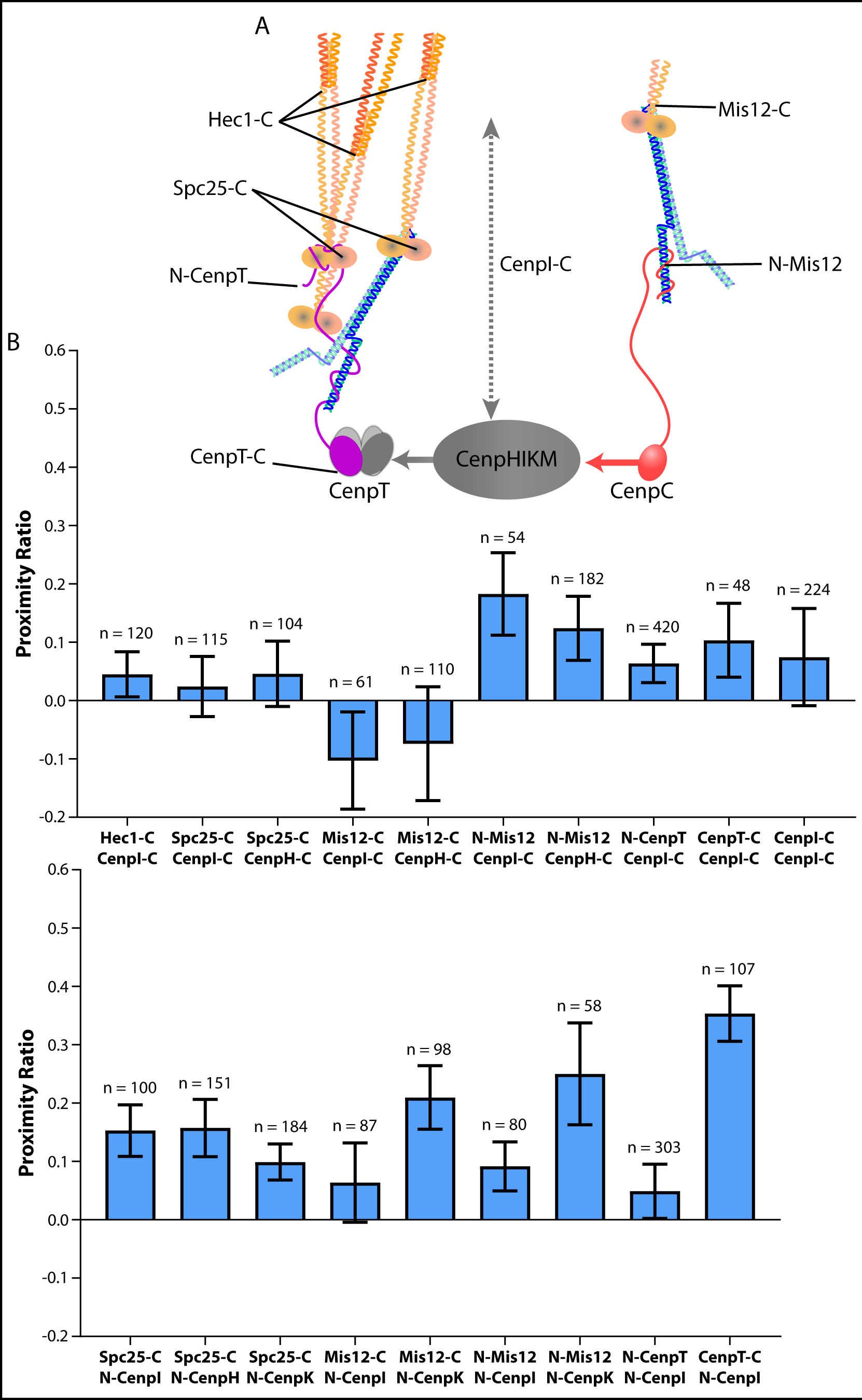
The N-termini, but not the C-termini, of members of the CenpHIKM complex produce weak FRET with Ndc80C and Mis12C. A) Diagram of the biochemical recruitment pathway for Ndc80C. The CenpHIKM bridges an interaction between CenpT and CenpC. Due to limited structural data, the location of the N- and C-termini for members of the CenpHIKM complex are unknown. Based on fluorescence co-localization data, the CenpI C-terminus is drawn as potentially extending toward the centromeric end of Ndc80C (gray, dashed arrow). B) FRET does not occur between the C-termini of either CenpI or CenpH and members of the outer kinetochore, CenpT, or between neighboring CenpI C-termini. C) Weak FRET occurs between the N-termini of CenpI, CenpH, and CenpK and members of the outer kinetochore. Moderate to strong FRET is produced between the N-terminus of CenpI and the C-terminus of CenpT, consistent with the known biochemical interaction between these two complexes. Data in B) and C) are the average proximity ratio ± S.E.M of fully occupied, metaphase kinetochores. The number of measurements is indicated above each bar.

**Figure S6.**
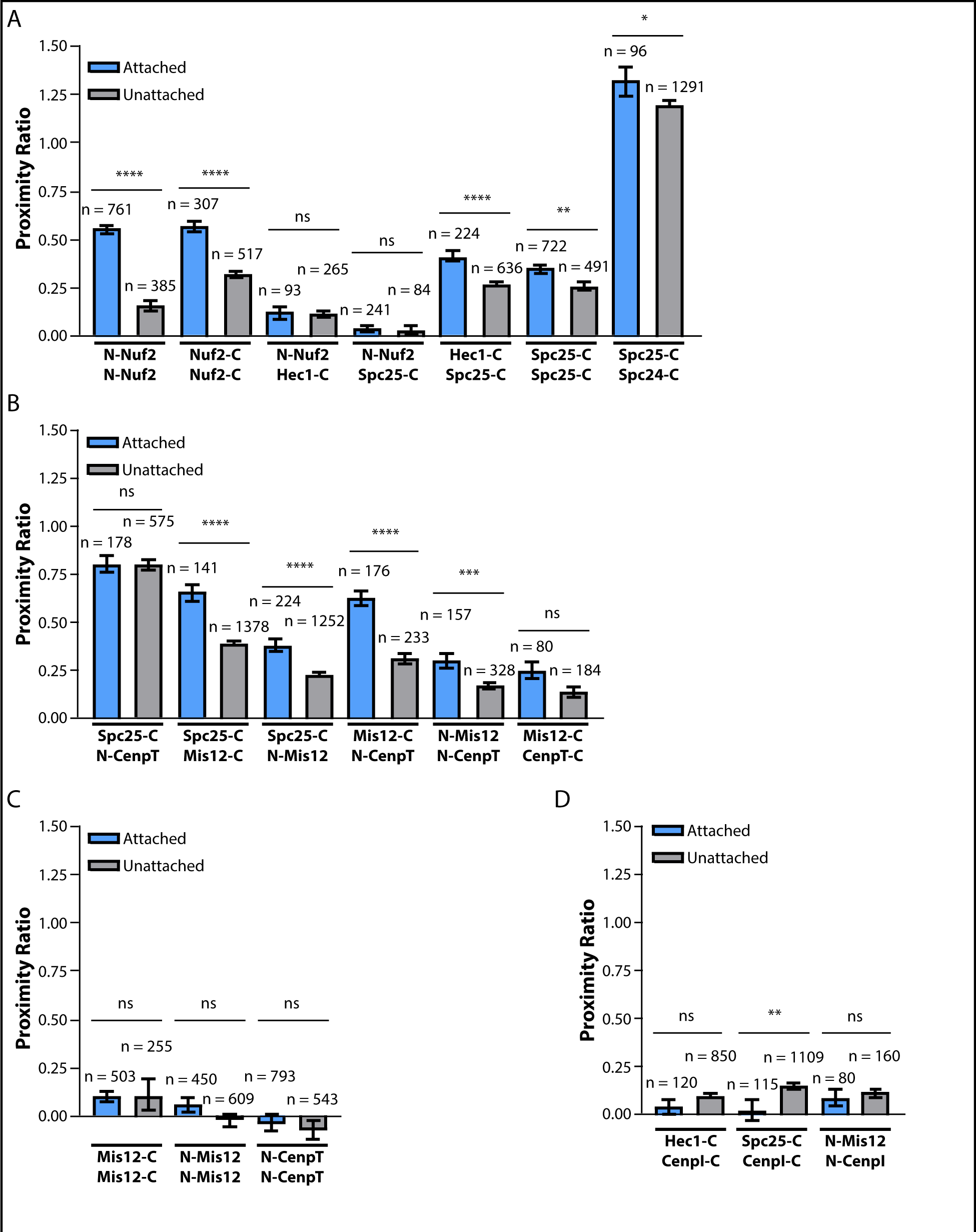
Additional FRET measurements of HeLa cells treated with nocodazole. A) – D) Bar graphs display the average proximity ratio ± S.E.M for the indicated FRET pairs. Light blue bars are measurements from metaphase aligned kinetochores and gray bars are measurements from nocodazole-treated, unaligned kinetochores. The number of measurements from fully occupied kinetochores is displayed above the bars for members of Ndc80C (A), proteins involved in Ndc80C recruitment (B), members of Mis12C (C), and between the outer kinetochore and CenpI (D). Statistical significance was evaluated by a Mann Whitney test, ns = not significant; * = p<0.05; ** = p<0.01; *** = p<0.001; **** = p<0.0001.

**Figure S7.**
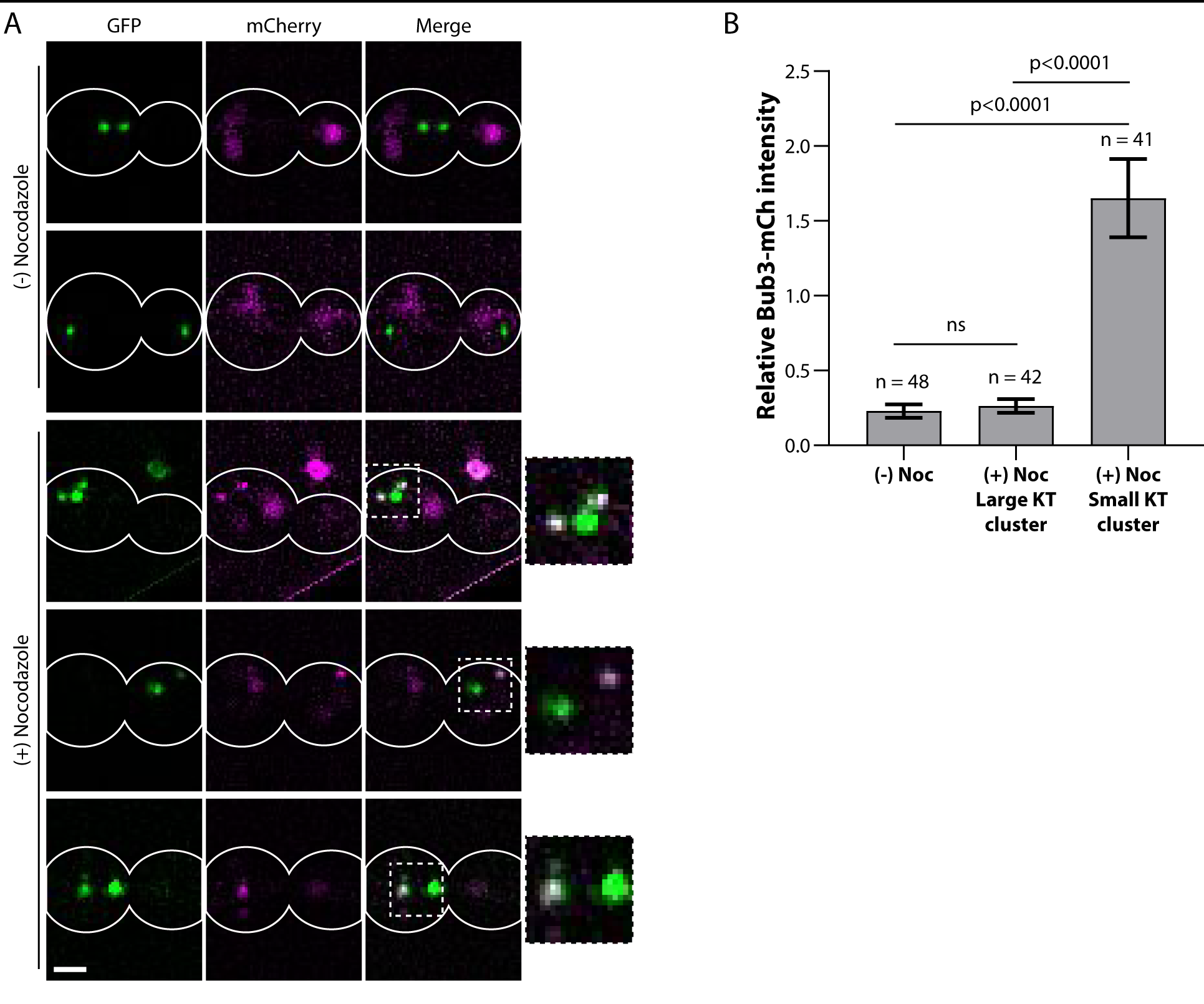
Nocodazole-treated budding yeast cells contain large clusters of microtubule-attached kinetochores and smaller clusters of unattached kinetochores. A) Example micrographs of budding yeast cells expressing Spc105-GFP and Bub3-mCherry, with or without the addition of nocodazole. White outlines indicate the cell membrane. Boxed regions in the merged images are shown as enlarged insets. Scale bar, 2 µm. B) The amount of Bub3-mCherry relative to Spc105-GFP was measured for metaphase kinetochores, as well as for the large kinetochore and small kinetochore clusters in nocodazole-treated cells. Small kinetochore clusters recruit higher amounts of Bub3, indicating the lack of microtubule attachments in this population.

**Figure S8.**
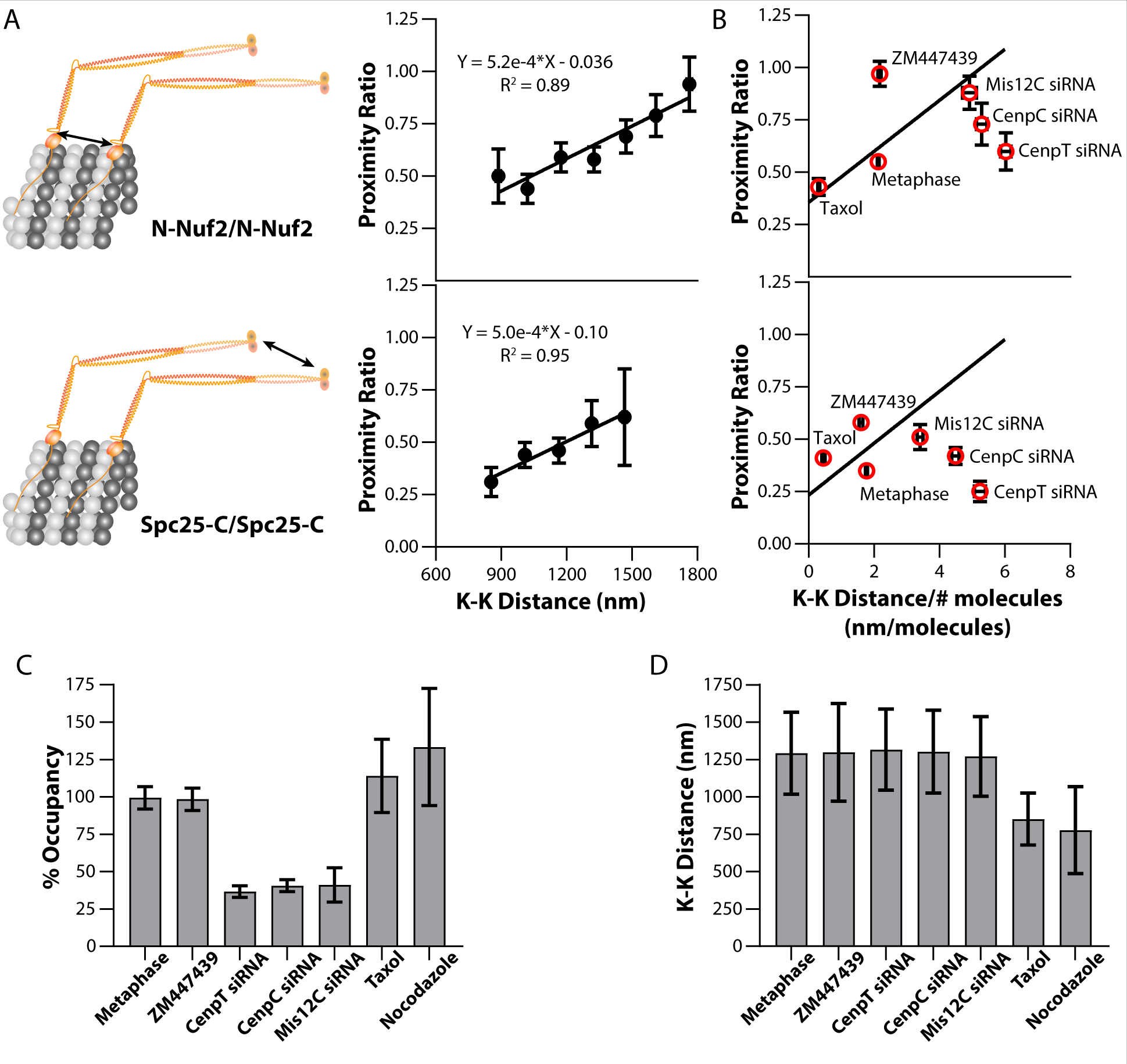
Relationship between the proximity ratio and the centromeric tension per molecule for Ndc80C with various drug and siRNA treatments. A) Cartoons for N-Nuf2/N-Nuf2 and Spc25-C/Spc25-C inter-complex FRET (left) and plots of the corresponding metaphase proximity ratios versus K-K distance (right, same as plots in Fig. 5A). B) The linear regressions in A) were re-scaled by dividing the sister kinetochore distance by the average number of molecules in a metaphase kinetochore. Plotted alongside the regressions are the average proximity ratios for kinetochores after various treatments, including: untreated metaphase, Taxol-treated, Aurora B kinase inhibition (ZM447439), CenpT siRNA, CenpC siRNA, and Mis12C siRNA treated kinetochores. C) The % occupancy and D) the average sister kinetochore separation distance (both relative to untreated metaphase kinetochores) is plotted for N-Nuf2/N-Nuf2 expressing HeLa cells after the indicated drug or siRNA treatments. Error bars in A) and B) are ± S.E.M. Error bars in C) and D) are ± S.D

**Figure S9.**
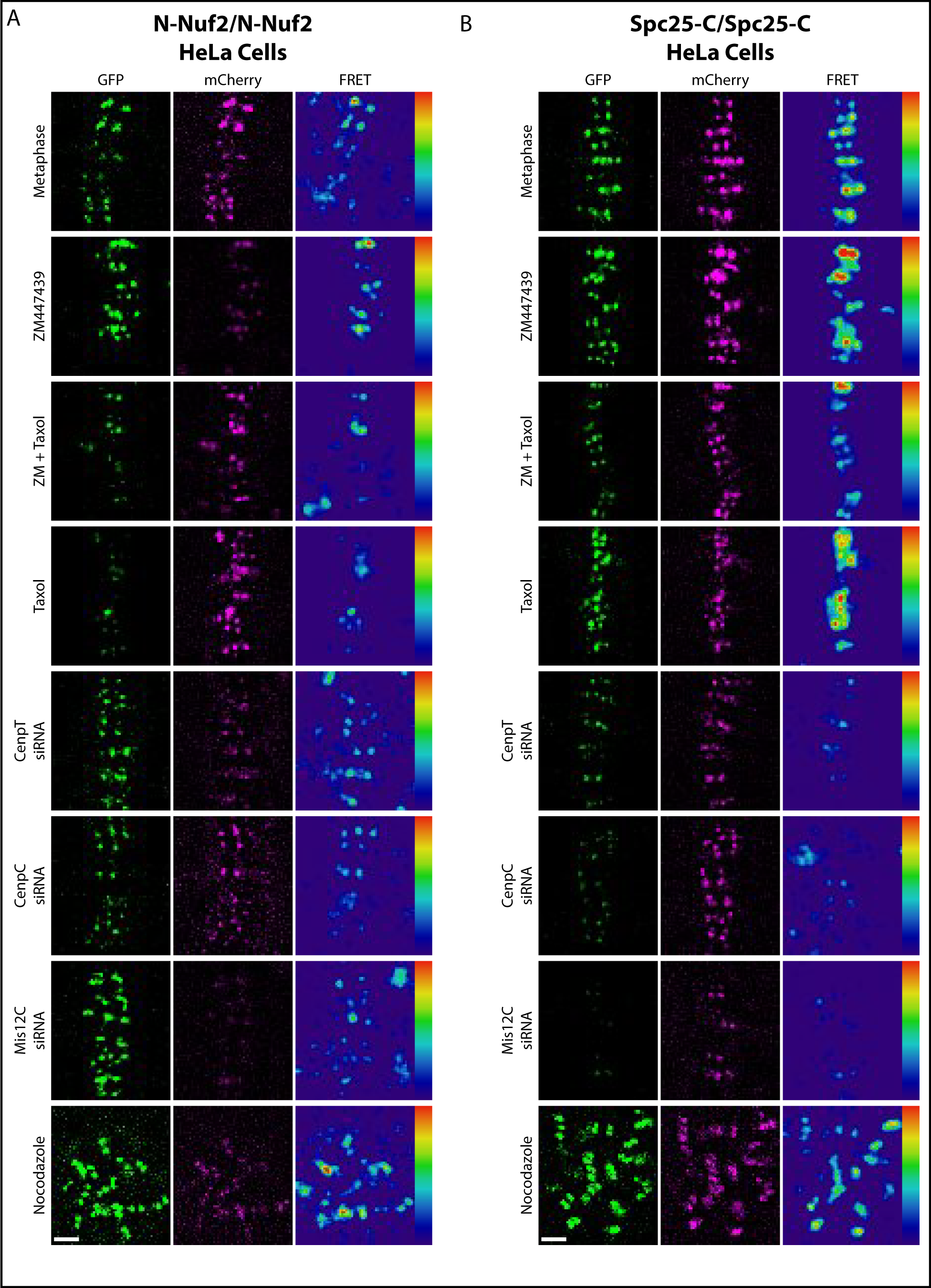
Example micrographs of HeLa cells after drug and siRNA treatments. A) GFP, mCherry, and FRET channel for N-Nuf2/N-Nuf2 expressing cells in the indicated treatments. Each channel is scaled equivalently for comparison between treatments. B) GFP, mCherry, and FRET channel for Spc25-C/Spc25-C expressing cells in the indicated treatments. Each channel is scaled equivalently for comparison between treatments. Scale bars, 2 µm. Note that images between the two cell lines are scaled differently.

